# Pathogenic *PTCD1* variants cause mitochondrial protein aggregation and cardiomyopathy

**DOI:** 10.64898/2026.06.06.730566

**Authors:** Ana Andjelković, Noah Wetterich, Michael Bull, Arthur Fox, Filippo Scialò, Daniella H. Hock, Benjamin B.A. Raymond, Langping He, Nichola Z. Lax, Matthias Trost, Zofia M. A. Chrzanowska-Lightowlers, Robert N. Lightowlers, Alberto Sanz, David A. Stroud, Jarno Mäkelä, Robert W. Taylor, Uwe Richter, Monika Oláhová

## Abstract

Disorders of mitochondrial oxidative phosphorylation affecting multiple respiratory chain complexes are among the most common causes of mitochondrial disease in humans. However, impaired energy metabolism alone does not fully account for tissue-specific vulnerability and disease progression, suggesting that additional molecular mechanisms contribute to disease pathology. We previously identified *PTCD1* variants in a child with infantile cardiomyopathy associated with a combined respiratory chain deficiency. Here, we establish the pathogenicity of three *PTCD1* (NM_015545.4) variants in which p.(Arg113Trp) and p.(Gly184Arg) segregate in *cis* whereas p.(Arg130*) is present in *trans,* demonstrating that disrupted mitochondrial proteostasis contributes to tissue damage in PTCD1 deficiency. PTCD1 patient cardiac tissue characterisation revealed impaired mitoribosome biogenesis, alongside increased aggregation of selective mitochondrial matrix proteins. Cell models expressing individual and combined *PTCD1* missense variants, coupled with proteomics, recapitulated the protein aggregation, with the *cis* p.(Arg113Trp);p.(Gly184Arg) combination showing the most severe effect. Protein aggregation was accompanied by altered OPA1 processing and mitochondrial network remodelling. Our findings establish accumulating proteotoxic stress arising from impaired mitoribosome assembly as a pathogenic mechanism in post-mitotic tissues, driving PTCD1 cardiomyopathy.

## Introduction

Pathological variants in protein components of the mitochondrial gene expression machinery often cause a combined biochemical defect in oxidative phosphorylation (OXPHOS), leading to a clinically and genetically heterogeneous group of rare multisystemic disorders (Boczonadi *et al*, 2018; Richter-Dennerlein *et al*, 2026; Taylor *et al*, 2014). The underlying pathogenic mechanisms are largely attributed to respiratory chain deficiency resulting from impaired mitochondrial gene expression and/or OXPHOS complex biogenesis. However, the marked tissue-specificity and progressive clinical course suggest that additional molecular mechanisms beyond the bioenergetic deficit exist, contributing to disease pathogenesis (Suomalainen & Battersby, 2018).

Biogenesis of the OXPHOS machinery requires effective coordination between nuclear and mitochondrial gene expression to facilitate translation of the 13 mitochondrial DNA-encoded OXPHOS subunits by the mitochondrial ribosome (Kramer *et al*, 2023; McShane & Churchman, 2024; Tang *et al*, 2020). The human mitoribosome (55S) consists of the 28S small subunit (mtSSU), which contains the 12S rRNA and 30 mitoribosomal proteins (MRPs), and the 39S large subunit (mtLSU), which contains the 16S rRNA together with 52 MRPs (Amunts *et al*, 2015; Itoh *et al*, 2021). Mitoribosomal biogenesis requires precise post-transcriptional modifications and processing of the 12S and 16S rRNAs, alongside coordinated assembly of the 82 nuclear-encoded MRPs. Several RNA granule components and mitochondrial auxiliary assembly factors have been identified, ensuring the formation of mitoribosomes (Brischigliaro *et al*, 2024).

Previous studies in mice identified pentatricopeptide repeat domain protein 1 (PTCD1) as an essential factor for 16S rRNA stability and mitoribosomal LSU biogenesis, functioning in a complex with FASTKD2 and RPUSD4 proteins in mitochondrial RNA granules (Antonicka & Shoubridge, 2015; Perks *et al*, 2017; Perks *et al*, 2018; Popow *et al*, 2015). Intriguingly, an Alzheimer’s disease (AD) association study identified the p.(Arg113Trp) *PTCD1* (NM_015545.4) variant as a low-penetrance risk allele enriched in AD patients (Fleck *et al*, 2019; Pa *et al*, 2019). This observation raises the possibility that defects in PTCD1 may contribute to neurodegeneration through molecular mechanisms that are beyond OXPHOS deficiency and are associated with imbalanced protein homeostasis.

Accordingly, defects in mitoribosomal biogenesis can lead to accumulation of unassembled MRP intermediates and formation of insoluble protein aggregates in the mitochondrial matrix. Dedicated safeguarding mechanisms have evolved to mitigate such damage in the form of mitochondrial protein quality control machinery (Collier *et al*, 2023; Ng *et al*, 2021; Richter *et al*, 2019). These include mitochondrial chaperones HSP60 (Bross & Fernandez-Guerra, 2016), mtHSP70 (Havalova *et al*, 2021) and TRAP1 (Joshi *et al*, 2022), alongside matrix proteases LONP1 (Zurita Rendon & Shoubridge, 2018) and CLPXP (Szczepanowska & Trifunovic, 2022). In addition to degrading damaged proteins, LONP1 also has important roles in RNA processing and the mitoribosome assembly itself (Zurita Rendon & Shoubridge, 2018).

When protein misfolding and/or aggregation exceeds the capacity of the mitochondrial protein quality control system, the resulting imbalance in mitochondrial matrix proteostasis can trigger proteotoxic stress, further impairing mitochondrial function. Pathogenic variants in *LONP1* cause cerebral, ocular, dental, auricular and skeletal syndrome (CODAS) (Strauss *et al*, 2015). Variants in *CLPP* are associated with Perrault syndrome, which presents with sensorineural hearing loss and primary ovarian insufficiency in females (Brodie *et al*, 2018; Jenkinson *et al*, 2013; Key *et al*, 2021), while variants in *HSPD1*, encoding HSP60, cause hereditary spastic paraplegia (Comert *et al*, 2020; Hansen *et al*, 2002). Together, this demonstrates that proteotoxicity in the mitochondrial matrix is sufficient to cause severe tissue-specific disorders. Proteotoxic stress poses a particular threat to post-mitotic cells such as neurons and cardiomyocytes, which lack the capacity to clear protein aggregates through cell division and are therefore at higher risk of progressively accumulating proteotoxic damage over time.

Here, we characterise the molecular basis for fatal infantile hypertrophic cardiomyopathy caused by segregating *PTCD1* variants. Using post-mitotic patient cardiac tissue and proliferating cell models, we establish how impaired mitoribosomal biogenesis drives proteotoxic stress, collapse of the mitochondrial network and combined OXPHOS deficiency in the heart. Together, our findings reinforce emerging evidence that mitochondrial proteostasis and network dynamics are key determinants of cellular health, whose disruption causes tissue damage independently of OXPHOS dysfunction, while OXPHOS impairment arises as a downstream consequence.

## Results

### *PTCD1* variants are associated with severe cardiomyopathy and combined OXPHOS deficiencies

We previously reported candidate variants in *PTCD1*, encoding an RNA-binding protein involved in mitochondrial ribosome biogenesis (Fleck *et al*., 2019; Perks *et al*., 2018; Schild *et al*, 2014), in a female infant who died of mitochondrial cardiomyopathy aged 8 months. This case was identified within a large cohort of patients with combined OXPHOS deficiencies who were subjected to gene-agnostic whole exome sequencing (Taylor *et al*., 2014). Sequencing identified three heterozygous variants in *PTCD1* (NM_015545.4): c.337C>T, p.(Arg113Trp); c.388C>T, p.(Arg130*) and c.550G>A, p.(Gly184Arg) (**Fig. 1A-B**). The c.337C>T, p.(Arg113Trp) *PTCD1* missense variant (*rs35556439*) is relatively common in the general population (MAF 2.937x10^-3^, with 7436 heterozygotes and 32 homozygotes in gnomAD v4) and has been robustly linked with AD in association studies (Fleck *et al*., 2019), where it acts as a low-penetrance risk allele. The *PTCD1* p.(Arg113Trp) residue shows low evolutionary conservation and high population frequency, indicating it is most likely not pathogenic in isolation. In contrast, the c.550G>A, p.(Gly184Arg) missense variant affects a highly conserved glycine residue located in the pentatricopeptide repeat (PPR) domain, a critical RNA-binding region, and is very rare (MAF 2.478x10^−5^, with 40 heterozygotes and no homozygotes in gnomAD v4). *In silico* analysis predicted the *PTCD1* c.550G>A, p.(Gly184Arg) variant to be "probably damaging" by PolyPhen-2, with a CADD score of 24.3 and REVEL score of 0.667. The c.388C>T, p.(Arg130*) variant is rare (MAF 5.824x10^−5^, with 94 heterozygotes and no homozygotes in gnomAD v4); and it is predicted to introduce a premature stop codon and is therefore anticipated to be subject to nonsense-mediated mRNA decay. Although other frameshift and nonsense *PTCD1* variants have been reported in gnomAD, none are common, and no loss-of function variants have been observed in the homozygous state, consistent with *PTCD1* being intolerant to bi-allelic loss-of-function variants.

**Figure 1.**
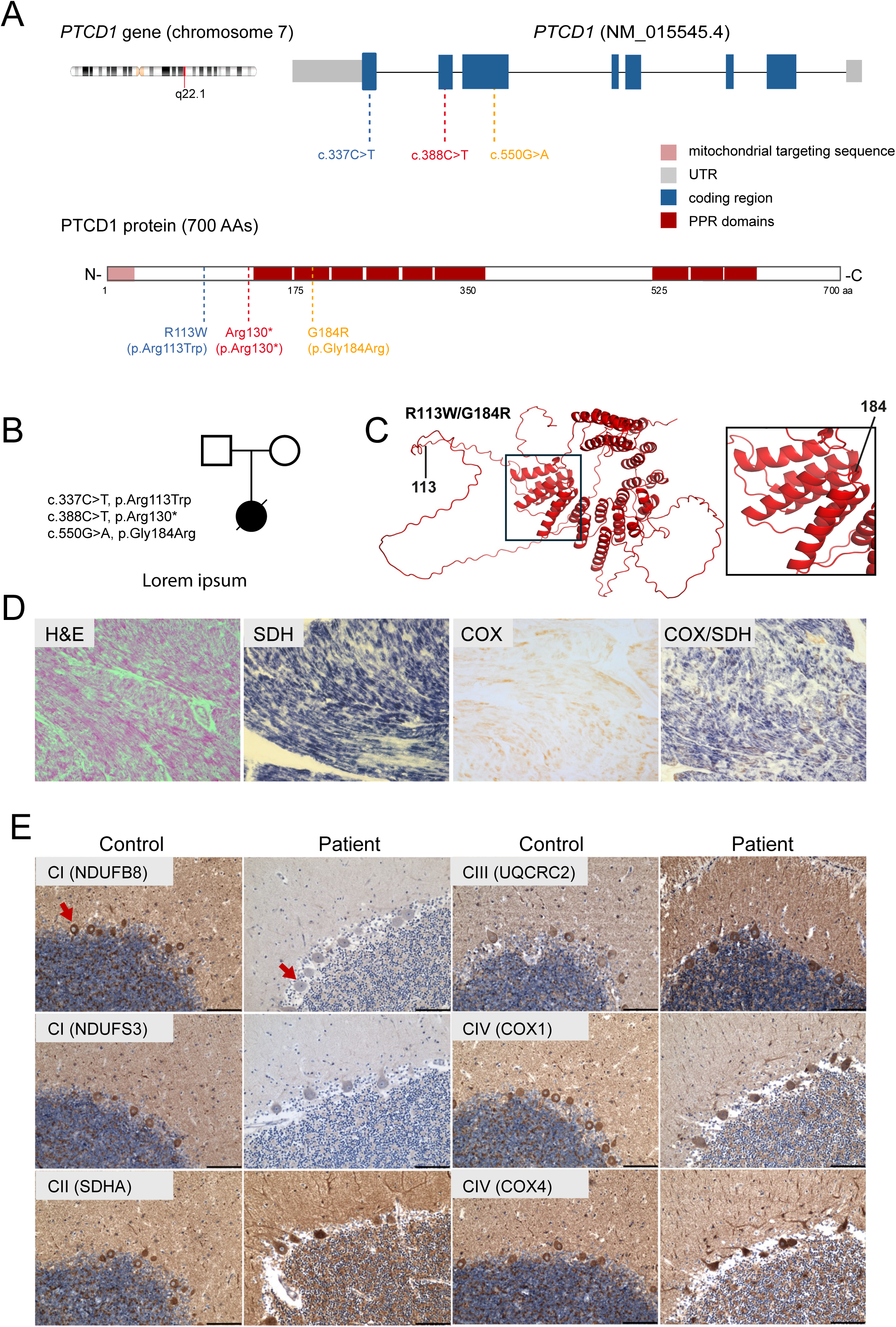
(**A**) A graphical representation of the *PTCD1* gene structure, illustrating the position of the mutated residues within *PTCD1* and the affected protein domains. (**B**) Family pedigree of the proband harbouring *PTDC1* variants. (**C**) *In silico* structural modelling of the human PTCD1 protein and variants using AlphaFold. (**D**) Mitochondrial respiratory chain deficiency in post-mortem cardiac muscle of the PTCD1 patient. (**i**) Haematoxylin & Eosin staining (**ii**) succinate dehydrogenase (SDH) (**iii**) cytochrome c oxidase (COX) and (iv) combined COX/SDH histochemistry (scale bar, 100 µm). (**E**) Mitochondrial respiratory chain deficiencies in post-mortem cerebellar tissue of the PTCD1 patient. Immunohistochemical staining of components of the mitochondrial respiratory chain revealed severe loss of complex I, as indicated by decreased NDUFB8 and NDUFS3 immunoreactivity in the Purkinje neurons indicated by red arrows. Immunoreactivity for the complex II subunit SDHA, complex III subunit UQCRC2 and mitochondria-encoded complex IV subunit I (COX1) was preserved, whilst the nuclear-encoded complex IV subunit IV (COX4) appeared slightly increased. Serial sections were stained with with haematoxylin, and the densely packed area of blue stained neurons represents the granule cell layer (scale bar, 100 µm).

No parental samples were available for segregation studies to determine the phase of the three identified *PTCD1* variants. We therefore isolated RNA from patient-derived cardiac tissue and performed allelic phasing studies. Sanger sequencing revealed that the c.337C>T, p.(Arg113Trp) and c.550G>A, p.(Gly184Arg) variants segregate together on one allele, while the c.388C>T, p.(Arg130*) variant is present in *trans* (**Fig. EV1**). This phasing, together with *in silico* pathogenicity predictions, indicates that the patient is compound heterozygous, carrying a complex allele, possibly hypomorphic, p.(Arg113Trp); p.(Gly184Arg) in *cis* with the predicted loss-of-function allele p.(Arg130*) in *trans*, resulting in impaired PTCD1 function. Overall, allele phasing indicates that the patient is compound heterozygous, carrying a hypomorphic complex allele along with a loss-of-function allele. In the absence of parental samples, we cannot exclude the possibility that the p.(Arg130*) variant arose *de novo;* however, the absence of clinical phenotypes in the parents suggests that haploinsufficiency alone is unlikely to be disease-causing.

PTCD1 contains PPR motifs that mediate RNA binding across all domains of life. As a protein-RNA chaperone, it also harbours intrinsically unstructured regions that are likely required for protein-protein interactions essential for its role in mitoribosome biogenesis. Structural modelling with AlphaFold (Jumper *et al*, 2021) predicts that the AD-associated p.(Arg113Trp) variant resides within an unstructured region, whereas the p.(Gly184Arg) variant localises to the PPR domain, important for the maintenance of helix-turn-helix repeats necessary for RNA recognition (**Fig. 1A and C**).

Diagnostic histochemical analysis of post-mortem cardiac tissue from the PTCD1 patient demonstrated a severe mitochondrial respiratory chain defect. Haematoxylin and Eosin (H&E) staining was employed to determine cardiac tissue morphology. Although limited by the image resolution, the overall tissue architecture was maintained (**Fig. 1D**). While succinate dehydrogenase (SDH) activity was preserved, consistent with intact mitochondrial content, cytochrome *c* oxidase (COX) staining demonstrated a marked decrease in enzymatic activity (**Fig. 1D**). Combined COX/SDH histochemistry further highlighted the predominance of COX-deficient/SDH-positive cardiomyocytes (**Fig. 1D**), indicative of a profound complex IV deficiency. In addition, immunohistochemical staining of mitochondrial respiratory chain components in fixed, post-mortem cerebellar tissue revealed severe deficiencies of nuclear-encoded complex I subunits NDUFB8 and NDUFS3 (**Fig. 1E**). Accredited diagnostic analyses of the patient’s mitochondrial genome (mtDNA), showed no evidence of mtDNA rearrangements, copy number abnormalities, or presence of likely-pathogenic mtDNA single nucleotide variants in the cardiac tissue.

We determined the activity of mitochondrial respiratory chain enzymes in the patient’s cardiac muscle homogenate and observed impaired activities across multiple respiratory chain enzymes (**Fig. 2A**). This is correlated with decreased steady-state levels of OXPHOS subunits, as analysed by immunoblotting (**Fig. 2B**). The patient’s cardiac muscle showed an almost complete loss of the mitochondrial-encoded complex IV subunits COX1 and COX2. We also observed a marked decrease in the nuclear-encoded complex I subunit NDUFB8 and complex III subunit UQCRC2 compared to controls (**Fig. 2B**). In contrast, the levels of nuclear-encoded complex II subunits SDHA and SDHB as well as the complex V subunit ATP5A remained relatively unchanged (**Fig. 2B**). The assembly of mitochondrial OXPHOS complexes was subsequently analysed by blue-native PAGE in control and PTCD1 patient cardiac tissue. Blue-native PAGE analysis showed markedly decreased amounts of fully assembled complexes I, I+III_2_, IV and V in PTCD1 patient cardiac muscle compared to controls, whereas the assembly profile of complex II was unaffected (**Fig. 2C**).

**Figure 2.**
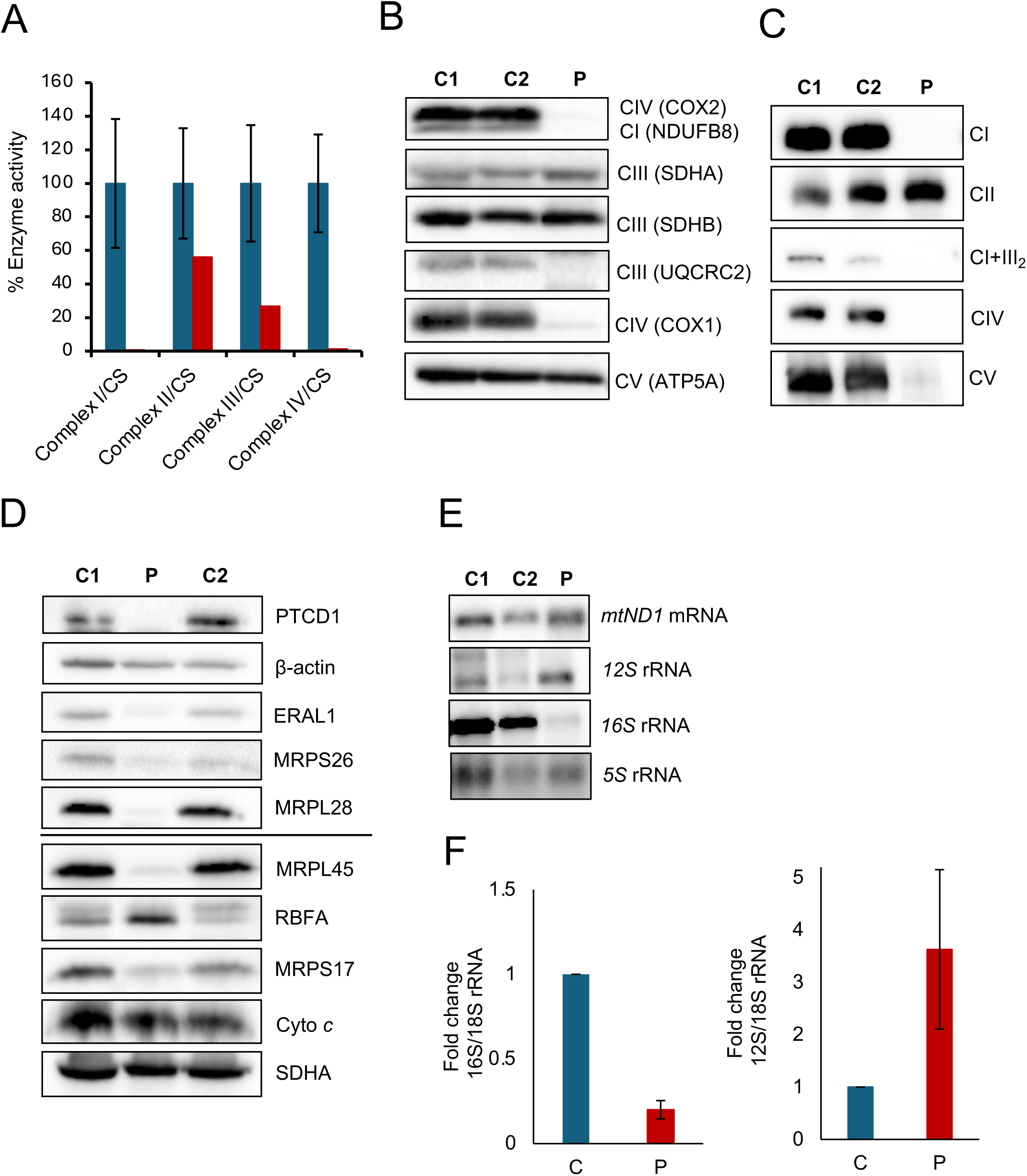
(**A**) Respiratory chain enzyme activities (CI–CIV) in mitochondria isolated from cardiac muscle of control (red) and PTCD1 patient (blue) with OXPHOS activities normalised to the citrate synthase activity. Mean enzyme activities for muscle controls (n = 25) are set at 100%. (**B**) Immunoblotting analysis of control (C1, C2) and PTCD1 patient (P) cardiac muscle lysates showing decreased steady-state levels of Complex I subunit NDUFB8, Complex III subunit UQCRC2 and Complex IV subunit COX2. Complex II subunit SDHB was used as loading control. (**C**) Blue-Native PAGE analysis showing an OXPHOS assembly defect in cardiac muscle from the PTCD1 patient, affecting the assembly of complexes I (NDUFA9), III (UQCRC2), IV (COX4) and V (ATP5A). Complex II (SDHA) was used as loading control. (**D**) Steady-state levels of mitoribosomal small subunit (mtSSU) and large subunit (mtLSU) proteins determined by immunoblotting in cardiac muscle lysates from controls (C1, C2) and PTCD1 patient. The levels of multiple mtSSU and mtLSU proteins and PTCD1 were decreased in the patient compared to control. SDHA and β-actin were used as loading controls. (**E**) Steady-state levels of mitochondrial 12S and 16S rRNAs and mt-ND1 mRNA were analysed by Northern blotting in the PTCD1 patient’s cardiac muscle, demonstrating a marked decrease in 16S rRNA. (**F**) Quantitative RT-PCR analysis showing decreased levels of 16S rRNA and increased levels of 12S rRNA in RNA isolated from PTCD1 patient’s cardiac muscle, normalised to 18S rRNA.

Consistent with these findings, analysis of the OXPHOS function in a *Drosophila melanogaster* model of PTDC1 deficiency demonstrated developmental defects and impaired mobility in eclosed flies, including failure of wing expansion (**Fig. EV 2A**). We have demonstrated that *PTCD1* knockdown in flies causes a significant decrease in oxygen consumption, measured by high-resolution respirometry (**Fig. EV 2B**). The findings in flies are consistent with the multiple respiratory chain deficiency observed in PTCD1 patient cardiac tissue, supporting the pathogenicity of PTCD1 and highlighting evolutionary conservation of its essential role across different species.

### *PTCD1* variants impair mitoribosome biogenesis in patient cardiac tissue

We next examined steady-state PTCD1 levels in soluble protein extracts from the patient’s cardiac muscle by SDS-PAGE and immunoblotting. PTCD1 protein levels were absent in the patient sample compared with age-matched controls (**Fig. 2D**), suggesting that the *PTCD1* variants adversely affect protein stability. PTCD1 function has been linked to biogenesis of the mitochondrial large ribosomal subunit in heart and skeletal muscle specific *Ptcd1* conditional knockout mice (Perks *et al*., 2018). To determine whether *PTCD1* variants underlie similar defects in human cardiac tissue, we examined the mitochondrial ribosome subunits in the soluble protein extracts by immunoblotting. We identified a marked decrease in multiple mitoribosomal proteins including MRPS26, MRPL28, MRPL45 and MRPS17 in the PTCD1 patient compared to controls. In addition, we found altered steady-state levels for two mitochondrial RNA-binding proteins: the human ribosome binding factor A (RBFA), which plays a crucial role in mitochondrial ribosome biogenesis (Rozanska *et al*, 2017) and the Era G-protein-like 1 (ERAL1) chaperone, which is required for mtSSU (12S rRNA) integrity (Dennerlein *et al*, 2010; Uchiumi *et al*, 2010) (**Fig. 2D**). This indicates that impaired PTCD1 function may disrupt the mitoribosome biogenesis machinery and not large subunit maturation alone. Next, we investigated the effect of *PTCD1* variants on the stability of mitochondrial rRNA transcripts in cardiac muscle samples from the PTCD1 patient and controls. Northern blotting analysis detected dramatically decreased steady-state levels of mitochondrial 16S rRNA transcripts (**Fig. 2E**), which was consistent with qRT-PCR data also showing a decrease in 16S rRNA (**Fig. 2F**). In contrast, the levels of 12S rRNA increased as determined by both Northern blot analysis and qRT-PCR (**Fig. 2E-F**). These data suggest that the *PTCD1* variants affect mitoribosomal maturation and assembly, indicating that a mitochondrial ribosome biogenesis defect likely impairs OXPHOS complex assembly (**Fig. 2C**). Importantly, our observations in PTCD1 patient cardiac tissue are entirely consistent with the mitoribosome defect observed in the cardiac-specific *Ptcd1* knockout mice (Perks *et al*., 2018), suggesting that *PTCD1* variants impair 16S rRNA processing and/or stability. Collectively, our data suggest that PTCD1 dysfunction prevents the proper maturation of mitoribosomes, supporting PTCD1’s role in mitoribosome biogenesis in humans.

### Cell models of *PTCD1* variants

As a patient-derived cell line was unavailable for further analysis, we investigated the cellular effects of the *PTCD1* variants on mitochondrial function using PTCD1-deficient cells described in (Fleck et al., 2019). The PTCD1 knockout cell line was transduced with either empty vector, wild-type *PTCD1,* or *PTCD1* variant constructs illustrated in **Figure 3A**. The variants p.(Arg113Trp) and p.(Gly184Arg), hereafter referred to as R113W and G184R, respectively, were expressed either individually or in combination as the *cis* p.(Arg113Trp); p.(Gly184Arg) variant, hereafter referred to as R113W/G184R (**Fig. 3A**).

**Figure 3.**
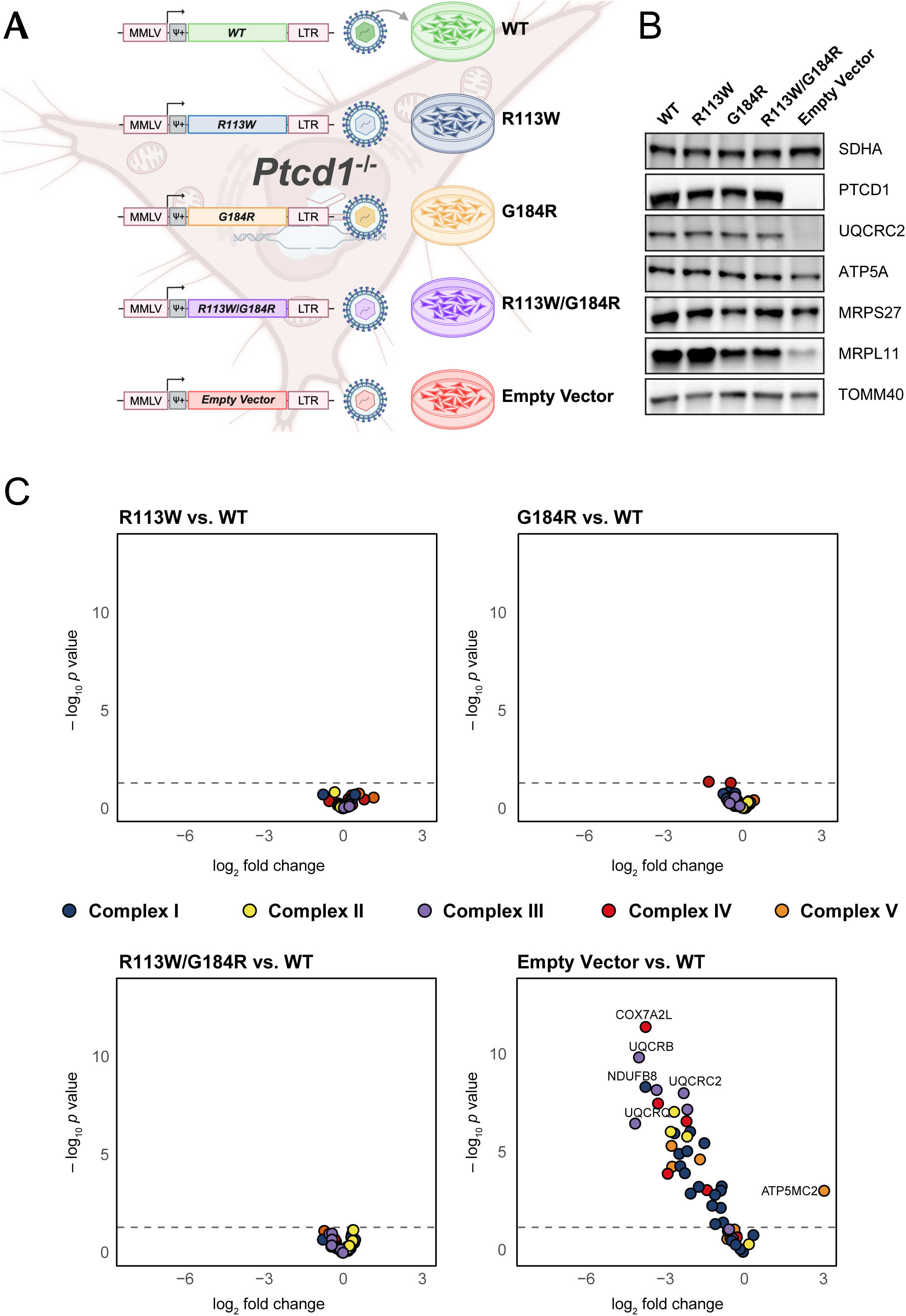
(**A**) Schematic representation of the *PTCD1* knockout–rescue strategy and the panel of *PTCD1* constructs introduced into *PTCD1-*deficient cells, with expression of empty vector, wild-type *PTCD1*, single missense variants p.(Arg113Trp) and p.(Gly184Arg), and the *cis* p.(Arg113Trp); p.(Gly184Arg) variant, indicated in the figure as R113W, G184R and R113W/G184R, respectively. (**B**) Immunoblotting analysis of mitochondrial respiratory chain subunits in *PTCD1* wild-type, knockout and variant HeLa cell lines. While empty vector transduced *PTCD1* knockout cells show decreased levels of ATP5A (complex V) and UQCRC2 (complex III), variant expression restored UQCRC2 and ATP5A to wild-type levels. Loss of PTCD1 in the knockout line is accompanied by a marked decrease in MRPL11, whereas expression of the wild-type and variants p.(Arg113Trp), p.(Gly184Arg) and p.(Arg113Trp); p.(Gly184Arg), shown as R113W, G184R and R113W/G184R, respectively, restores MRPL11 abundance close to wild-type levels. MRPS27 levels remain largely unchanged across all conditions. The nuclear encoded complex II subunit SDHA was used as loading control. (**C**) Quantitative LC–MS/MS whole cell proteomic analysis showing log2 fold change in the abundance of complexes I, II, III, IV and V subunits relative to wild-type for each variant cell line. Statistical significance was calculated by *t*-test in limma (*n* = 4). Respiratory chain subunits are restored to wild-type levels in all *PTCD1* expressing variant cell lines.

Our analysis of patient heart tissue suggested that variant PTCD1 proteins may disrupt mitoribosome biogenesis. To determine which individual or combined PTCD1 missense variants affected this process, we assessed steady-state levels of mitoribosomal proteins in the variant cell lines by immunoblotting. Solubilised protein fractions from PTCD1 wild-type, variants, and the empty vector PTCD1 knockout confirmed complete loss of PTCD1 protein in the knockout and robust expression in all transduced cell lines, with minor variation (**Fig. 3B**). In line with previous reports (Fleck *et al*., 2019; Perks *et al*., 2018) mtSSU MRPS27 protein levels remained largely unchanged across all cell lines (**Fig. 3B**). This supported the interpretation of PTCD1 as an mtLSU biogenesis factor that does not significantly influence the mtSSU maturation. In contrast, the protein steady-state level of the mtLSU subunit MRPL11 was markedly decreased in the *PTCD1* knockout but not severely decreased in cells expressing G184R and R113W/G184R *PTCD1* variant proteins (**Fig. 3B**). MRPL11 levels were unchanged when compared to wild-type in cells expressing the R113W variant (**Fig. 3B**).

Impaired mtLSU assembly capacity would be expected to hinder the translation of mtDNA-encoded respiratory chain subunits and, consequently, to disrupt the assembly of complexes I, III, IV and V. Expression of wild-type *PTCD1* and each of the *PTCD1* variants restored the respiratory chain complex defect (**Fig. 3B**), with ATP synthase subunit 5A levels restored from partial depletion and the complex III subunit UQCRC2 from complete loss compared to the *PTCD1* knockout cell line (**Fig. 3B**). The recovery of complex III and ATP synthase subunit levels is concordant with the minor decrease in mtLSU steady-state abundance observed in the cell lines expressing G184R alone and R113W/G184R (**Fig. 3B**).

Given the semi-quantitative nature of immunoblotting and to detect alterations in steady-state levels of mitoribosomal and respiratory chain proteins quantitatively, we performed whole-cell proteomic analysis using LC–MS/MS. The loss of PTCD1 resulted in the expected severe decrease in respiratory chain complexes I, III, IV and V (**Fig. 3C**). The knockout cell line transduced with empty vector also revealed the loss of some complex II proteins. This is noteworthy since complex II is purely nuclear-encoded. The abundance of the respiratory chain proteins from all respiratory chain complexes was fully restored in the cell lines expressing *PTCD1* variant proteins including R113W/G184R (**Fig. 3C**). However, we observed a coordinated decrease of mtSSU and mtLSU proteins not only in the *PTCD1* knockout line but likewise in the cell lines expressing G184R alone and R113W/G184R (**Fig. 4**). The R113W-expressing cell line showed no significant deviations relative to the wild-type, whereas the number of significant differentially expressed mitoribosomal proteins increased progressively in the order: G184W < R113W/G184R < *PTCD1* knockout (**Fig. 4**). This phenotype is most likely attributable to the inability of the G184W and of R113W/G184R cells to support normal mitoribosome biogenesis, mirroring a stepwise increase in severity from G184R to R113W/G184R to *PTCD1* knockout. Taken together, these findings indicate that both, the complete loss of PTCD1 and the dysfunction caused by *PTCD1* variants, lead to an overall depletion in mitoribosome abundance in proliferating cells, albeit with severity modulated by the R113W variant when in *cis* with G184R. While the R113W allele can rescue mitochondrial large subunit biogenesis, the G184R variant alone and R113W/G184R show progressively diminished capacity to maintain mtLSU abundance. These findings are consistent with a role for PTCD1 in mtLSU biogenesis and suggest that the *cis* combination of R113W and G184R variants is associated with reduced mtLSU assembly capacity, while OXPHOS function surprisingly appears comparatively preserved.

**Figure 4.**
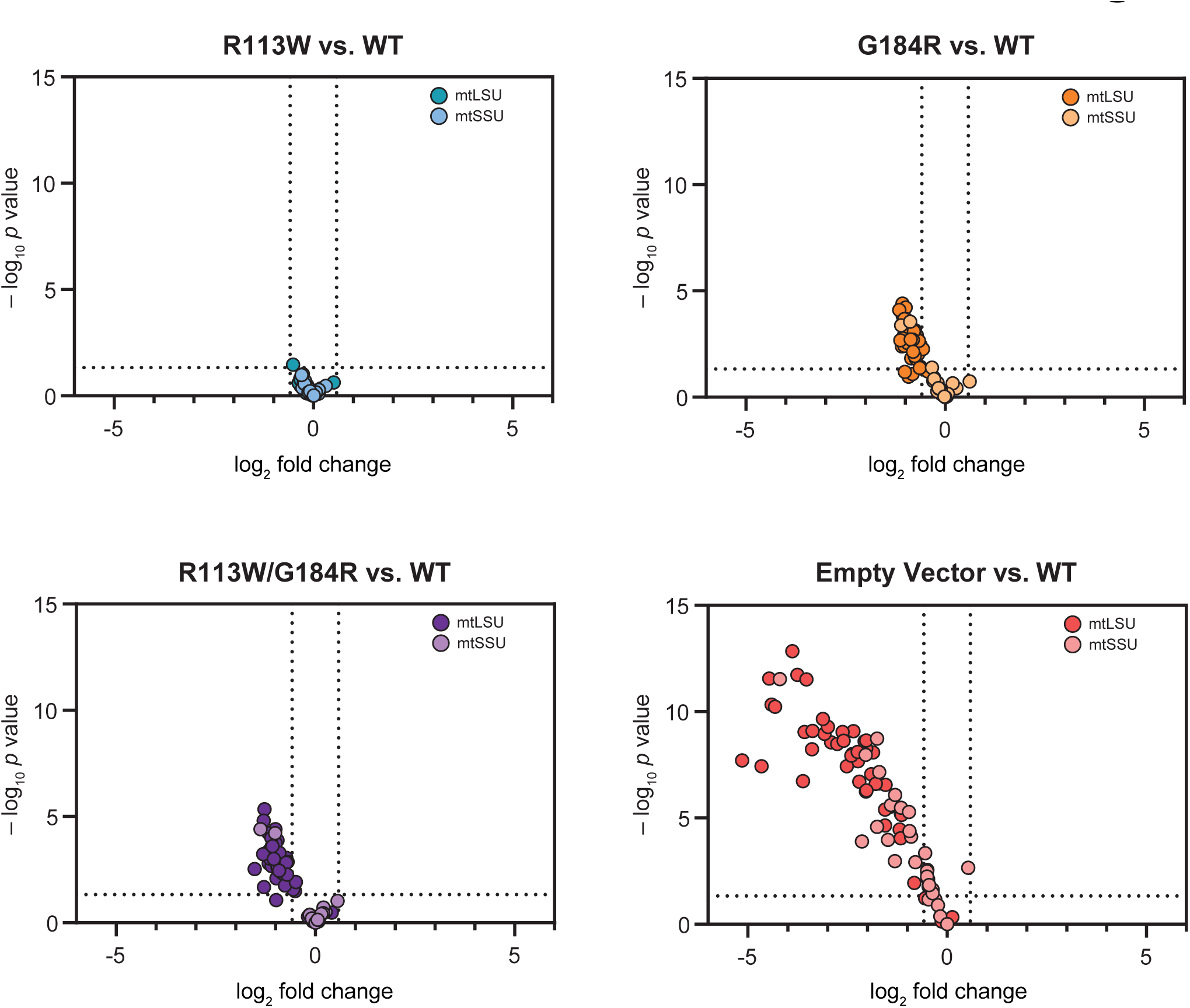
*PTCD1* variants differentially affect mitoribosome biogenesis in cell models. Quantitative LC–MS/MS proteomic analysis of whole-cell lysates from *PTCD1* wild-type, knockout and variant-expressing lines showing log_2_ fold change of mitochondrial ribosomal proteins relative to wild-type. MtSSU and mtLSU proteins are displayed separately, revealing a coordinated decrease of both subunits in the knockout and a stepwise increase in deficiency more pronounced for mtLSU proteins in the order p.(Gly184Arg) < p.(Arg113Trp); p.(Gly184Arg) < *PTCD1* knockout, while the p.(Arg113Trp) variant doesn’t show a significant deviation from wild-type. Statistical significance was calculated by *t*-test in limma (*n* = 4).

### Mitochondrial protein aggregation in patient heart tissue and variant cell lines

While the reconstituted cell lines revealed decreased mitoribosome abundance with comparatively modest effects on OXPHOS function, immunoblotting analysis of patient heart tissue indicated an apparent decrease in both mitoribosomal and OXPHOS subunits, prompting us to perform a comprehensive mitochondrial proteome analysis of patient heart tissue. We performed LC–MS/MS analysis on isolated cardiac mitochondria from the PTCD1 patient and age-matched controls. Consistent with previous immunoblotting analysis (**Fig. 2B**) and *PTCD1* knockout cell line proteomics (**Fig. 3C**), we observed a decrease in multiple OXPHOS subunits, affecting complexes I, III and IV (**Fig. 5A-B**). In addition, the levels of multiple mitochondrial RNA granules associated proteins and processing/modification factors were increased in the PTCD1 patient cardiac tissue (**Fig. 5A**). Mitochondrial proteomic analysis of MRPs revealed an unexpected expression pattern of mtLSU and mtSSU proteins. The abundance of mtLSU proteins was relatively normal in the PTCD1 patient compared to control, whereas the mtSSU proteins were more abundant in the patient cardiac tissue (**Fig. 5C**). To visualise the spatial distribution of these changes, we mapped the log_2_ fold change abundances onto the mitoribosome cryo-EM structure (PDB: 3J9M), which clearly shows the asymmetric pattern of mtLSU preservation and strong increase in abundance of mtSSU proteins across the mitoribosome structure (**Fig. 5D**). Rather than observing a clear decrease in MRPs in the patient cardiac tissue homogenates, the relative preservation of MRPs in the proteomic dataset (**Fig. 5C**) may reflect extraction under the harsher denaturing conditions used for LC–MS/MS sample preparation, consistent with both mtLSU and mtSSU proteins being less soluble under the milder extraction conditions used for immunoblotting (**Fig. 2D**). To test this, we isolated soluble and insoluble protein fractions from control and PTCD1 patient cardiac tissue homogenates using a sequential detergent-based extraction method, where the soluble fraction was isolated following sonication and RIPA buffer lysis, and the insoluble fraction was resolubilised by sonication in RIPA buffer containing 2% SDS. In line with our proteomic data, we found selective accumulation of the mtSSU MRPS27 and mtLSU MRPL28 in the insoluble fraction of the patient sample by immunoblotting (**Fig. 5E**). This was accompanied by co-sedimentation of other mitochondrial matrix proteins involved in mitochondrial gene expression, including the matrix protease LONP1, RNA processing factor MRPP3 and PTCD1 itself. In contrast, when we assessed the redistribution of proteins, the intermembrane space protease HTRA2 and the inner membrane translocase TIMM50 were found to be absent from the insoluble fraction, suggesting compartment specificity. Consistent with the decrease in complex I subunit NDUFB8 detected by immunoblotting (**Fig. 2B**), we observed decreased levels of NDUFB8 in both soluble and insoluble fractions. In contrast, the complex II subunit SDHA was absent only from the insoluble fraction of the patient samples (**Fig. 5E**). The levels of the TCA cycle enzyme citrate synthase, which also resides in the mitochondrial matrix showed similar levels between patient and controls in the insoluble fraction, suggesting that there may be selective precipitation of mitoribosome-associated proteins, rather than generalised matrix protein aggregation. In addition, immunoblotting of protein lysates from control and PTCD1 cardiac tissue used for proteomic analysis, which uses harsher lysis conditions, confirmed that the levels of MRPL28 were comparable to controls, whilst MRPS17 levels were increased and the complex I subunit NDUFB8 decreased in the PTCD1 patient (**Fig. EV4**). We found the levels of the mitochondrial matrix chaperone HSP70 and outer mitochondrial membrane protein FIS1 similar in the PTCD1 patient compared to controls, further supporting selective redistribution of the mitoribosomal proteins in PTCD1 cardiac tissue (**Fig. EV4**). Together, our findings demonstrate that MRPs are not absent in PTCD1 patient heart tissue, but instead accumulate in an insoluble aggregated state, consistent with the increased vulnerability of post-mitotic tissue to proteotoxic stress.

**Figure 5.**
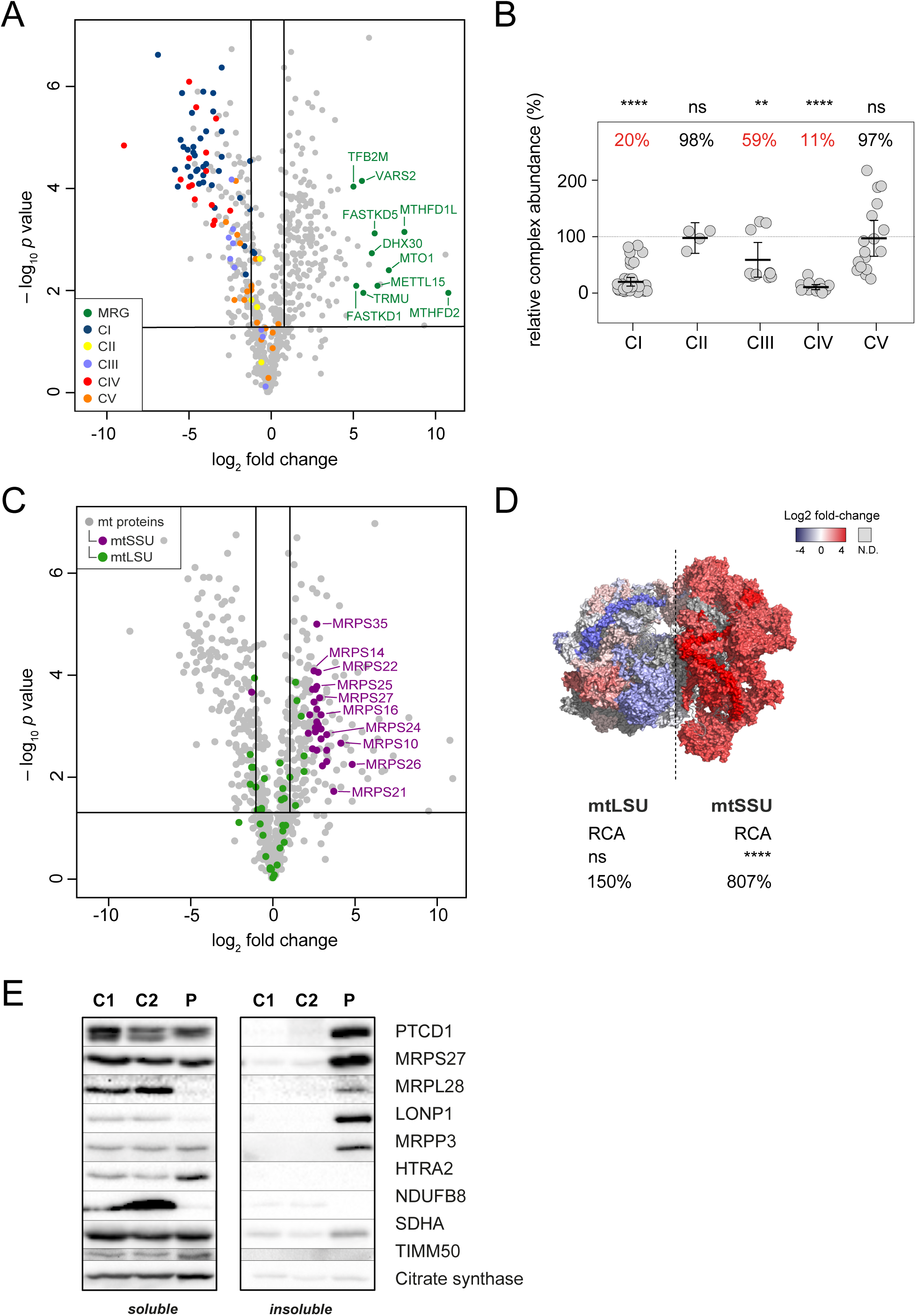
(**A**) Quantitative LC-MS/MS proteomic analysis of mitochondria isolated from cardiac tissue of controls and PTCD1 patient showing a global decrease in subunits of complexes I, III and IV. The volcano plot shows the mean log_2_ fold change and the −log_10_ of the two-tailed Student’s *t*-test. *p* value of proteins quantified in mitochondria isolated from PTCD1 patient’s cardiac tissue compared to age-matched controls. Individual OXPHOS complexes (CI-V) and mitochondrial RNA granule (MRG) proteins are colour coded. Horizontal line represents p = 0.05 and vertical lines represent log_2_ fold-change ± 1. (**B**) Relative complex abundance of OXPHOS complexes in the PTCD1 patient cardiac mitochondria relative to controls. Each data point indicates an individual subunit, the middle bar represents mean complex abundance, whilst the upper and lower bars represent 95% confidence interval. Statistical significance was calculated using a two-sided *t*-test between the individual protein means **** p < 0.0001, ** p < 0.01, ns = not significant, p > 0.05.(**C**) Volcano plot of cardiac mitochondrial proteins highlighting the large and small mitoribosome subunits (mtLSU and mtSSU) from PTCD1 patient compared to controls, showing an increase in mtSSU subunits. Horizontal line represents p = 0.05 and vertical lines represent log_2_ fold change ± 1. (**D**) Log_2_ fold change abundances of mtLSU and mtSSU from two-tailed Student’s *t*-test mapped onto the mitoribosome cryo-EM structure (PDB: 3J9M). RCA = relative complex abundance analysis. N.D = not detected. **** p < 0.0001, ns = not significant, p > 0.05. (**E**) Immunoblotting analysis of the soluble and insoluble fractions of controls (C1, C2) and PTCD1 patient cardiac muscle (P) showing a selective accumulation of mitochondrial ribosomal proteins in the insoluble fraction of the PTCD1 patient compared to controls. SDHA and citrate synthase serve as compartment specific controls for the inner membrane associated fraction and matrix solubility, respectively.

To assess putative protein aggregation in the variant cell lines, insoluble protein fractions (pellet fractions) obtained during cell lysis and protein solubilisation in the above experiments were subjected to LC–MS/MS analysis. To control for changes in overall protein abundance and quantify mitoribosomal subunit aggregation associated with PTCD1 dysfunction or loss, we derived an “aggregation score” for each protein defined as the log2 fold change in the insoluble fraction minus the log_2_ fold change in the soluble fraction (**Figs. 5 and 6A**). These aggregation scores were visualised as a heat map to compare the aggregation profiles across all cell lines (**Fig. 6B**). The PTCD1 R113W expressing cell line consistently exhibited the lowest aggregation scores among the variants for both small and large subunit proteins (**Fig. 6A-B**), whereas the PTCD1 G184R expressing cell line showed higher scores, particularly for mtLSU components (**Fig. 6B**). The PTCD1 R113W/G184R expressing cells displayed the highest mtLSU aggregation scores among the variant cell lines, while the *PTCD1* knockout (empty vector) showed the greatest overall mitoribosomal protein aggregation, again most pronounced for mtLSU proteins (**Fig. 6B**). Collectively, these data define an allelic series in which complete loss of PTCD1 causes maximal mitoribosomal protein aggregation, R113W exerts the mildest effect, and acts as a modifier when in *cis* with G184R, resulting in a more severe aggregation phenotype. Although, aggregation of mitoribosomal proteins was clearly detectable in proliferating R113W/G184R variant cells, this phenotype is substantially more pronounced in post-mitotic cardiac tissue.

**Figure 6.**
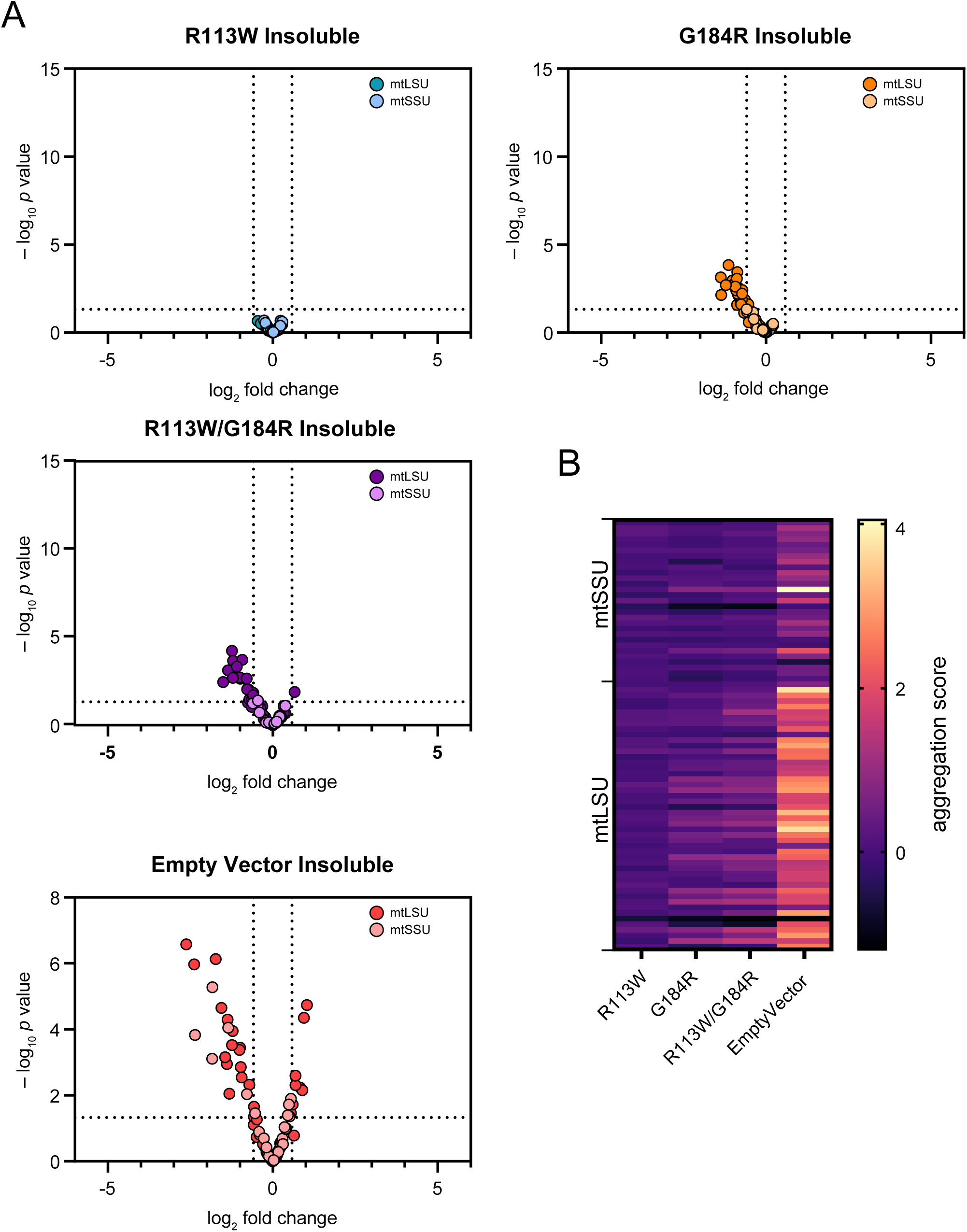
PTCD1 dysfunction drives mitoribosomal protein aggregation. (**A**) Volcano plots depicting log_2_ fold change comparisons of mitoribosomal proteins from variant cell lines relative to wildtype. Statistical significance is indicated as −log_10_ *p* value, with the significance threshold set at of p = 0.05. Proteins of mtLSU and mtSSU are colour coded. In the *PTCD1* knockout and the R113W/G184R variant, the number of mitochondrial ribosomal proteins that were not significantly decreased is higher in the insoluble fraction compared to the soluble (see **Fig. 4**). (**B**) Heat map of aggregation scores for individual proteins across the *PTCD1* variant cell lines and the knockout. mtLSU and mtSSU proteins are shown as separate blocks in the heat map allowing comparison of aggregation patterns between the two mitoribosomal subunits, with individual proteins arranged by row and the different cell lines in columns. The colour scale indicates the aggregation score for each protein, calculated as the difference between the log_2_ fold change in the insoluble fraction and the log_2_ fold change in the soluble fraction (see **Fig. 4**).

### Mitochondrial stress response is activated in PTCD1 patient

The observed defect in mitoribosomal assembly with concomitant protein aggregation in the mitochondrial matrix prompted us to investigate how impaired mitoribosomal biogenesis and protein aggregation affect mitochondrial homeostasis in the PTCD1 patient post-mortem cardiac muscle. We examined OPA1 processing, as altered OPA1 cleavage is an indicator of mitochondrial stress (Anand *et al*, 2014; Baker *et al*, 2014). PTCD1 patient cardiac tissue showed altered OPA1 processing compared to control tissues (**Fig. 7A**). Loss of the long OPA1 (L) isoforms in the patient, together with relative accumulation of the shortest OPA1 isoform, S5, indicates proteolytic cleavage by the stress-activated inner mitochondrial membrane protease OMA1 in response to disturbed proteostasis in the mitochondrial matrix. Immunoblotting analysis of *PTCD1* knockout and variant cell lines show modest OMA1 activation in the *PTCD1* knockout cells, indicated by a decreased abundance of the short S4 isoform and a concomitant increase in the OMA1 cleavage-derived S3 and S5 isoforms (**Fig. 7B**). In contrast, expression of *PTCD1* variants R113W/G184R, did not alter the pattern of OPA1 isoforms (**Fig. 7B**), suggesting that these conditions do not exceed the threshold required to trigger an OMA1-dependent stress response.

**Figure 7.**
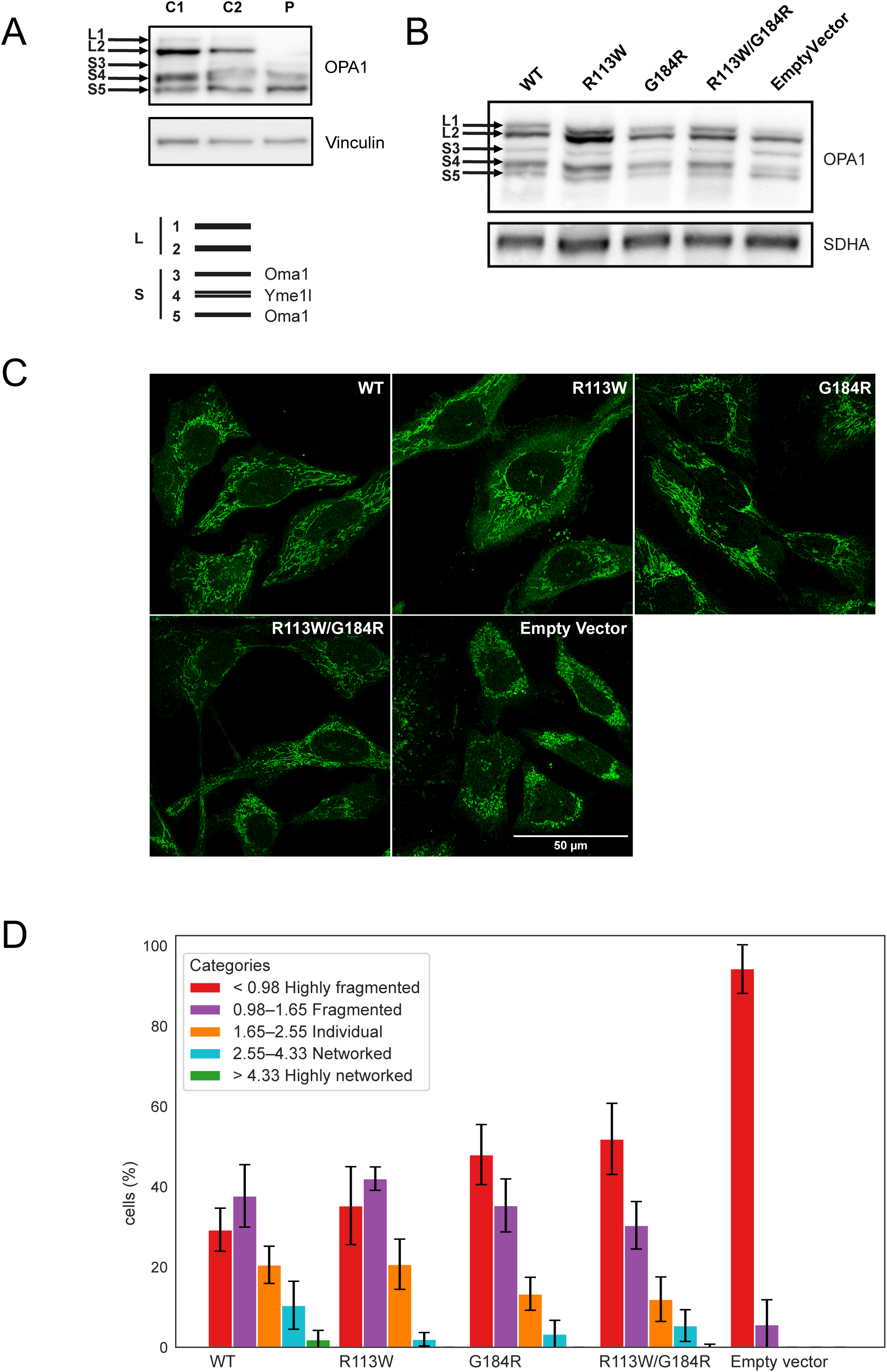
*PTCD1* variants induce mitochondrial stress signalling and mitochondrial network fragmentation. (**A**) Immunoblot analysis of cardiac muscle from the PTCD1 patient and age-matched controls showing altered OPA1 processing in the patient, characterised by loss of the long OPA1 (L) isoforms and accumulation of the shortest S5 isoform, indicating OMA1 activation. (**B**) OMA1-driven alteration of OPA1 processing, with relative accumulation of the S3 and S5 OMA1-derived cleavage products and loss of S4, was observed only in *PTCD1* knockout but not in the *PTCD1* variant cell lines. (**C**) Analysis of mitochondrial morphology based on fragment size in confocal microscopy. Categories were generated using the Gaussian mixture model (S4) and include all variant cell lines. The graph shows an elevation in fragmentation from the control through the variant and peaking in the knockout cells. Graphs include three independent experiments for statistical significance.

Cleavage of the membrane-anchored long OPA1 isoforms is associated with altered mitochondrial ultrastructure, and complete *PTCD1* knockout has been shown to cause mitochondrial fragmentation in both mouse tissue (Perks *et al*., 2018) and human cells (Fleck *et al*., 2019). To assess mitochondrial morphology in the variant cell lines, we analysed mitochondrial ultrastructure by confocal microscopy. This revealed an intermediate mitochondrial network fragmentation phenotype that became progressively more pronounced relative to wild type in the order: R113W < G184R < R113W/G184R < *PTCD1* knockout (**Fig. 7C-D**). Across all PTCD1 variant-expressing cell lines, we observed a systematic shift in mitochondrial network organisation, with increasing PTCD1 perturbation associated with a higher proportion of cells in the more severe fragmentation categories.

Fragmentation states were quantified by assigning individual cells to one of five discrete degrees of fragmentation (**Fig. 7C**). These categories were defined using a Gaussian mixture model fitted to the pooled mitochondrial fragment size data from all conditions, which generated four objective threshold gates at the intersections of the Gaussian components and thereby established a common fragmentation scale across the entire dataset. Using this classification, both single-mutant *PTCD1* lines already exhibited an increased representation of cells in intermediate-to-high fragmentation categories compared with control, whereas R113W/G184R and *PTCD1* knockout cells were markedly enriched in the highest fragmentation category, with a corresponding loss of cells with low fragmentation (**Fig. 7C**).

The variability and reproducibility of mitochondrial fragment size within each condition were further illustrated by plotting biological replicates separately, confirming that each genotype displayed a characteristic and internally consistent fragmentation profile across replicates (**Fig. EV5**). These quantitative findings were supported by qualitative confocal imaging of ATP5A-labelled mitochondria, which recapitulate the shift from predominantly tubular networks in control cells to progressively more punctate and fragmented mitochondria in the single variants, to R113W/G184R and *PTCD1* knockout cells. This pattern closely mirrored the redistribution of cells across fragmentation categories and the shift in fragment size distributions observed in the bar graph and violin plot analyses (**Fig. EV5**).

Taken together, these findings indicate that, although OXPHOS levels are fully rescued upon expression of any of the patient missense variants, mitochondrial fragmentation is observable in both knockout and R113W/G184R patient variant-expressing cell lines.

## Discussion

In this study, we report pathogenic variants in *PTCD1* that drive cardiomyopathy through a primary defect in mitoribosome biogenesis, accompanied by mitochondrial proteotoxicity and combined OXPHOS deficiency. By integrating analyses of post-mortem patient heart tissue with PTCD1-deficient and variant-reconstituted cell models, we show that defective mtLSU assembly likely leads to aggregation of mitochondrial gene expression proteins, activation of stress-responsive remodelling of the mitochondrial network, and, ultimately, combined respiratory chain deficiency, a phenotype that was also recapitulated in a *D. melanogaster* model of PTCD1 knockdown.

Our findings challenge the view that bioenergetic failure is the sole driver of OXPHOS disease and instead highlight proteotoxic stress arising from unassembled mitoribosomes, particularly in post-mitotic tissues. Patient cardiac tissue harbouring *PTCD1* variants p.(Arg113Trp); p.(Gly184Arg) in *cis* with p.(Arg130*) in *trans* (**Fig. EV6**) exhibits combined OXPHOS deficiencies, depleted soluble mitoribosomal proteins, and selective aggregation of matrix proteins including MRPS27 and MRPL28 mitoribosomal subunits, PTCD1, LONP1, and MRPP3. Molecular cloning of *PTCD1* patient cDNA confirmed the compound heterozygous phasing of the variants. However, in the absence of parental samples, we cannot exclude the possibility of *de novo* occurrence. Nevertheless, the healthy clinical presentation of both parents suggests an autosomal recessive mode of inheritance, indicating that a single pathogenic *PTCD1* variant is not sufficient for disease manifestation.

In cell models, expression of the individual missense *PTCD1* variants did not significantly alter steady-state levels of respiratory chain complexes, in contrast to the severe OXPHOS deficiency present in the patient cardiac tissue. However, the variant cell lines recapitulated an allelic series of defects, with the *cis* R113W/G184R variant causing the most severe mtLSU instability, protein aggregation, OPA1-mediated stress responses, and mitochondrial fragmentation. Interestingly, aberrant mitochondrial ultrastructure has been reported in the liver mitochondria of *Ptcd1^+/–^* mice, which exhibited mild uncoupling, decreased L-OPA1 levels, and increased S-OPA1 abundance, but only minimal changes in respiratory chain protein levels (Perks *et al*., 2017). Together, these findings emphasise that both OXPHOS deficiency and proteostasis collapse might be the drivers of pathology in patient heart tissue. Increased HTRA2 abundance in PTCD1-deficient cardiac tissue likely reflects a compensatory attempt to clear misfolded or aggregated mitochondrial proteins generated by impaired mitoribosome biogenesis and translation failure, in keeping with the established role of HTRA2 as a stress-inducible, aggregation-responsive mitochondrial protease (Gray *et al*, 2000; Toyama *et al*, 2022).

Consistent with previous findings (Fleck *et al*., 2019), the common c.337C>T, p.(Arg113Trp) variant (MAF 2.937x10^-3^), a low-penetrance Alzheimer’s risk factor, appears benign in isolation but exacerbates mtLSU instability and protein aggregation in *cis* with p.(Gly184Arg) in the variant cell lines. Notably, respiratory chain complexes accumulation did not significantly differ between the variant lines. Earlier work linking AD risk to energy deficits has suggested PTCD1 dysfunction as a potential indicator for mitochondria’s role in AD (Pa *et al*., 2019). Our findings extend this hypothesis by highlighting proteostasis failure as a primary consequence of variant PTCD1’s defective RNA-protein chaperoning activity, with energetic failure arising as a downstream consequence.

We observed a clear decrease in 16S rRNA, whereas mt-ND1 mRNA levels were increased, despite ND1 protein levels being significantly downregulated in the patient’s cardiac proteome. This pattern is consistent with a defect at the level of mitochondrial translation rather than transcription or RNA processing. In line with previous studies in cardiac-specific *Ptcd1^-/-^*knockout mice, our data support a role for PTCD1 in 16S rRNA maturation, which is essential for mtLSU biogenesis (Perks *et al*., 2018). In addition, cardiac mitochondrial proteomics revealed elevated steady state levels of several mitochondrial RNA processing factors, including the 16S rRNA maturation factor FASTKD2, which has been shown to associate with PTCD1 and RPUSD4 within the 16S rRNA complex, the 12S rRNA methyltransferase METTL15, and additional RNA processing factors including FASTKD1 and FASTKD5. The compensatory upregulation of these factors or, alternatively, their collapse into protein aggregates, may represent a direct consequence of the residual PTCD1 activity conferred by the p.(Arg113Trp); p.(Gly184Arg) allele, which remains insufficient to sustain 16S rRNA maturation and downstream mtLSU assembly. This observation may further indicate that chronic perturbation of mitochondrial RNA processing is recognised by the cell, inducing elevated expression of RNA processing factors, or alternatively reflect high sensitivity for maladaptive aggregation within mitochondrial RNA granules. Furthermore, the increase in 12S rRNA levels, despite decreased ERAL1 abundance, indicates a failure of mtSSU assembly, likely resulting in accumulation of unassembled rRNA. This is consistent with a broader disruption of mitoribosome biogenesis in PTCD1-deficient tissue, where defects in mtLSU maturation secondarily impair mtSSU stability and assembly.

In a recent study, mitochondrial proteostatic stress in human cells has been shown to induce protein aggregates within the mitochondrial matrix along with extensive cristae remodelling and altered processing of the cristae shaping OPA1 (Ehses *et al*, 2026), although the precise mechanistic sequence remains to be established. Molina-Riquelme *et al*. show that OPA1 downregulation, together with decreased cristae density observed in aged mice, precedes changes in mitochondrial morphology (Molina-Riquelme *et al*, 2026), representing an early hallmark of cardiac aging.

Our findings for proteins associated with mitochondrial aggregates in the patient heart are in line with recent findings which indicate that mitochondrial protein aggregation sequesters assembly factors and proteins of the small ribosomal subunit (Holthusen, 2025 bioRxiv). The results demonstrate that mitochondrial ribosome assembly is particularly susceptible to proteostatic stress in the mitochondrial matrix, offering a structural and mechanistic explanation for the accumulation of assembly factors, mitochondrial RNA-granule components, LONP1, and mtSSU proteins in the protein aggregates of the patient heart due to compromised mtLSU biogenesis.

More broadly, our findings also closely align with reports implicating mitochondrial proteostasis dysfunction in human disease. Variants in the major mitochondrial matrix chaperone HSP60 are associated with neurodegeneration such as spastic paraplegia (Hansen et al., 2002), whereas LONP1 variants cause CODAS syndrome and cause mitochondrial protein aggregation (Strauss et al., 2015). These observations extend and contextualise the findings here, where PTCD1 related mitoribosome instability is accompanied by selective aggregation of mtLSU proteins and matrix resident factors, together with OXPHOS deficiency and mitochondrial fragmentation.

Increased HTRA2 protease activity has been reported in the frontal cortex of patients with AD, where it has been proposed to represent a compensatory response to proteotoxic stress and accumulation of misfolded proteins (Westerlund *et al*, 2011). Consistent with this, the elevated HTRA2 levels observed in PTCD1 patient cardiac tissue likely reflect activation of mitochondrial quality control pathways in response to defective protein homeostasis. This complements the increased LONP1 and CLPX expression observed in the Perks model. Together, these findings suggest coordinated activation of multiple mitochondrial quality control systems in response to failed ribosome assembly.

These results underscore the vulnerability of the mitochondrial translation machinery and suggest that aggregation of mitochondrial ribosomal proteins represent a shared pathomechanism across mitochondrial diseases, and possibly other age-related disorders in which mitochondria are compromised, including AD. The apparent dissociation between rescue of steady-state OXPHOS protein abundance and persistence of mitochondrial fragmentation and aggregation phenotypes in the variant cell lines implies that restoration of respiratory chain subunits alone may not be sufficient to restore mitochondrial homeostasis once proteostatic imbalance has been initiated. This suggests that mitochondrial proteotoxicity itself may become an independent pathogenic driver downstream of impaired translation.

Our findings substantially extend the model proposed by Perks *et al*., who identified PTCD1 as a critical regulator of 16S rRNA maturation, mtLSU biogenesis, and mitochondrial translation. While impaired mitoribosome assembly in the *Ptcd1^-/-^* knockout model activated retrograde mTOR signalling and pro-survival transcriptional programmes, our data suggest that defective mtLSU assembly additionally triggers mitochondrial proteotoxic stress characterised by aggregation of mitochondrial gene expression factors, activation of mitochondrial quality control pathways, altered OPA1 processing, and mitochondrial fragmentation. Together, these observations support a model in which failed mitoribosome biogenesis initiates a coordinated stress response involving OMA1-dependent mitochondrial remodelling, protease activation, and mito-nuclear signalling pathways aimed at maintaining cellular survival despite severe mitochondrial dysfunction.

## Materials and Methods

### Ethics statement

Informed consent for diagnostic and research studies was obtained in accordance with the Declaration of Helsinki protocols and approved by the relevant ethical guidelines issued by the institutional ethics committees.

### Histological, histochemical and biochemical analyses

Post-mortem cardiac tissue was available in limited quantities from a single patient; all analyses were performed using the entirety of the available material." Postmortem tissue from the patient cardiac muscle was subjected to diagnostic histopathological analyses including cytochrome *c* oxidase (COX), succinate dehydrogenase (SDH), and sequential COX-SDH according to established protocols (Old & Johnson, 1989). Mitochondrial respiratory chain complexes and citrate synthase activities were determined spectrophotometrically in mitochondrial fractions isolated from control and patient cardiac muscle homogenates according to previously described methods (Kirby *et al*, 2007).

Immunohistochemical analyses were performed on age-matched control and patient cerebellar sections, which were immunostained with NDUFB8, NDUFS3, SDHA, UQCRC2, COXI and COXIV antibodies, using a polymer detection system (Menarini Diagnostics) and visualisation with 3,3′-diaminobenzidine (Sigma-Aldrich) as previously described (Lax *et al*, 2012). The availability of the post-mortem PTCD1 patient cardiac and brain tissue was limited, (n = 1).

### Immunoblotting of patient cardiac tissue

To obtain soluble protein fractions, cardiac tissues (∼10mg) were homogenised in liquid nitrogen using a pestle and mortar. Each sample was resuspended in 0.5 mL of RIPA buffer containing 1% Igepal CA-630, 1.5% Triton X-100, 0.5% sodium deoxycholate, 0.1% SDS, 10 mM β-mercaptoethanol, 1 mM phenylmethylsulfonyl fluoride (PMSF), and 1× Ethylenediaminetetraacetic acid (EDTA) free protease inhibitor cocktail (Pierce) and incubated on ice for 45 min. Following 2×15 sec homogenisation with a polytron homogeniser, lysates were cleared by centrifugation at 17 000 g for 10 min at 4°C and the soluble fraction was retained. A modified method was used to separate the soluble and insoluble proteins from cardiac tissues, which included a sonication step (10 sec on/off for a period of 3 min) prior to the 45 min incubation on ice. Following homogenization with polytron homogeniser as described above and centrifugation at 17 000 g for 10 min at 4°C, the soluble supernatant was retained. The resulting pellet representing the insoluble fraction was resuspended in RIPA buffer containing 2% SDS and sonicated using 10 sec on/off cycles for 1 min. The Pierce BCA Protein Assay Kit (Thermo Fisher Scientific) was used to determine protein concentrations. Equal amounts of protein from soluble and insoluble fractions were loaded onto SDS-PAGE gels and separated by electrophoresis using a Bio-Rad system, followed by a transfer of proteins on to a PVDF membrane (Immobilon-P, Millipore).

Immunoblotting was performed overnight at 4°C, unless otherwise indicated, using antibodies against PTCD1 (Abcam, ab121620, 1:500), NDUFB8 (Abcam, ab110242, 1:1000), SDHA (Abcam, ab14715, 1:1000), UQCRC2 (Abcam, ab14745, 1:1000), COX1 (Abcam, ab14705. 1:1000), ATP5A (Abcam, ab14748. 1:1000), Porin/VDAC1 (Abcam, ab14734, 1:2000), Total OXPHOS Human WB Antibody Cocktail (Abcam, ab110411, 1:1000), Vinculin (Abcam, ab219649, 1:2000), HTRA2 (R&D Systems, AF1458, 1:1000), β-actin (Sigma-Aldrich, A1978,1:10,000), OPA1 (BD Transduction Laboratories, 612606, 1:1000), MRPL28 (Proteintech, 21604-1-AP, 1:1000), MRPL45 (Proteintech, 15682-1-AP, 1:1000), MRPS17 (Proteintech, 18881-1-AP, 1:1000), MRPS26 (Proteintech, 15989-1-AP, 1:1000), MRPS27 (Proteintech, 17280-1-AP, 1:1000), LONP1 (Sigma-Aldrich, HPA002192, 1:500), MRPP3 (Abcam, ab185941, 1:1000), RBFA (Custom made Affinity purified, rabbit), ERAL1 (Proteintech, 11478-1-AP, 1:1000), TIMM50 (Proteintech, 22229-1-AP, 1:1000) and Citrate synthase (Abcam, ab96600, 1:5000). Species-specific HRP-conjugated secondary antibodies (Dako P0260, 1:2000 and P0399, 1:3000) were used for 1h at room temperature. Immunoblots were visualised using SuperSignal™ West Pico PLUS Chemiluminescent Substrate (Thermo Fisher Scientific) and imaged on a ChemiDoc MP Imaging System (Bio-Rad) using Image Lab software.

### Mitochondrial isolation and blue-native PAGE analysis of patient cardiac tissue

Crude mitochondrial extracts were isolated from cardiac tissues solubilised using n-dodecyl β-d-maltoside (DDM). OXPHOS complexes were separated by blue-native PAGE as described previously in (Olahova *et al*, 2017). Control and patient cardiac tissues (∼20 mg) were homogenised in a homogenization buffer containing 250 mM sucrose, 20 mM Imidazole-HCl pH 7.4, 1 mM PMSF, using a Teflon glass Dounce homogenizer at 4°C (20 strokes). Homogenised samples were centrifuged at 20 000 g for 10 min at 4°C and pellets washed 2x with 1 ml of homogenization buffer and centrifuged at 20 000 g for 5 min at 4°C. The final pellets were solubilised with DDM (Sigma-Aldrich) at 2 g/g protein (Hornig-Do *et al*, 2014) in 1X NativePAGE™ Sample Buffer (ThermoFisher Scientific) on ice for 20 min. Protein concentrations were determined with the Pierce BCA Protein Assay Kit (Thermo Fisher Scientific). Following centrifugation at 100 000 g for 15 min at 4°C the supernatants were retained, mixed with Coomassie G-250 (Sigma-Aldrich) 0.25% (w/v) and equal amount (50–100 μg) of control and patient samples was loaded onto NativePAGE™ Bis-Tris Mini Protein Gels, 4 to 16% (Thermo Fisher Scientific). OXPHOS complexes were electrophoretically separated using the X-Cell Sure Lock system (Thermo Fisher Scientific) and NativePAGE™ cathode and anode buffers (Thermo Fisher Scientific) at 4°C, followed by a wet transfer onto PVDF membranes (Immobilon-P, Millipore) and immunoblotting for CI (NDUFB8), CII (SDHA), CIII (UQCRC2), CIV (COXI) and CV (ATP5A) using primary and secondary antibodies described above.

### RNA extraction, Northern blot analysis and qRT-PCR

Total RNA was extracted from ∼20mg of age-matched control and patient cardiac tissues using ReliaPrep™ RNA Tissue Miniprep System (Promega) or TRIzol® (Invitrogen) according to manufacturer’s guidelines. RNA samples (1–3 µg) were denatured and electrophoretically separated on a 1.2% (w/v) agarose gel under denaturing conditions and subsequently transferred to a Genescreen Plus membrane (Life Science Products, Inc.). 32P-dCTP-radiolabelled probes were generated by random hexamer priming of PCR products corresponding to internal regions of 5S, 12S, 16S, and mt-ND1 genes. Typhoon FLA 9500 (GE/Fujifilm) was used to visualise the radiochemical signals. cDNA synthesis from RNA samples was carried out using the M-MLV Reverse Transcriptase (Promega) and following manufacturer’s instructions. Quantitative real-time PCR was performed using FastStart Essential DNA Green Master (Roche) ready-to-use reaction mix and the following primer pairs: 18S forward primer (GTAACCCGTTGAACCCCATT) and 18S reverse primer (CCATCCAATCGGTAGTAGCG), 12S forward primer (ACACTACGAGCCACAG) and 12S reverse primer (ACCTTGACCTAACGTC), 16S forward primer (CCAATTAAGAAAGCGTTCAAG) and 16S reverse primer (CATGCCTGTGTTGGGTTGACA).

### Cardiac mitochondrial proteomics

Mitochondria from controls and patient cardiac tissues were isolated following homogenization in 3ml of homogenization buffer containing 20 mM HEPES-KOH (pH 7.6), 220 mM Mannitol, 70 mM sucrose, 1 mM EDTA supplemented with containing freshly added 0.5 mM PMSF. Tissue homogenisation was performed using a glass/Teflon Potter homogeniser for approximately 50 strokes with a drill-mounted pestle. Mitochondria were separated by differential centrifugation for 5 min at 800 g at 4°C and the supernatant was transferred into a fresh tube and centrifuged for 10 min at 11 000 g at 4°C. The mitochondrial pellet was resuspended in 300 µl of homogenization buffer and the isolation was repeated as described above. The final mitochondrial pellet was resuspended in 100 µl of sucrose storage buffer. Pierce™ BCA Protein Assay Kit (Thermo Fisher Scientific) was used to determine protein concentrations in each sample. Fifty micrograms of isolated mitochondria was lysed in 50 µl lysis buffer containing 1% (w/v) SDC, 100 mM Tris pH 8.1, 40 mM 2-chloroacetamide (CAA) and 10mM Tris(2-carboxyethyl)phosphine hydrochloride (TCEP). Samples were vortexed and boiled for 15 min at 99°C with 1400 rpm shaking. Denatured lysates were allowed to cool to room temperature before adding 1:100 of trypsin (sample pH ∼8) and incubation at 37°C with overnight shaking. Samples were acidified with 100 µl ethyl acetate, 1% trifluoroacetic acid (TFA) for > 2 min at RT and loaded onto SDB-RPS (Supelco) stage tips, washed with 200 µl ethyl acetate, 1% TFA. Following another wash with 200 µl 0.2% TFA, proteins were eluted with 100 µl 80% acetonitrile (ACN), 1% ammonium hydroxide. Eluates were dried by vacuum centrifugation (SpeedVac) and peptides were reconstituted in 0.1% (v/v) TFA and 2% (v/v) ACN with brief sonication on water bath sonicator and analyzed by online nano-HPLC/electrospray ionisation-MS/MS on a Q Exactive Plus mass spectrometer connected to an Ultimate 3000 HPLC (Thermo Scientific) using a non-linear in-house 128 min gradient. (Cox & Mann, 2008; Hock *et al*, 2025; Hock *et al*, 2020; Rath *et al*, 2021; Tyanova *et al*, 2016)

### Drosophila melanogaster stocks and maintenance

*D. melanogaster* flies were cultured at 25°C on standard media containing 1% agar, 1.5% sucrose, 3% glucose, 3.5% dried yeast, 1.5% maize, 1% wheat, 1% soya, 3% treacle, 0.5% propionic acid, 0.1% Nipagin, with a controlled 12 h:12 h light:dark cycle. Knockdown of the *Drosophila PTCD1* orthologue (CG4611) was achieved using the UAS-GAL4 system. A line carrying a w1118; P[GD11371] v21898 construct against CG4611 was obtained from Vienna Drosophila Resource Center. Knockdown efficiency was confirmed by qPCR as described previously (Scialo *et al*, 2016b). Control flies carried the da-GAL4 driver without the UAS-RNAi transgene.

### *Drosophila melanogaster* high-resolution respirometry

Twenty whole male wild-type and *PTCD1* KD flies were homogenized in a homogenisation buffer containing 120 mM KCl, 5 mM KH_2_PO_4_, 3 mM Hepes, 1 mM EGTA, 1 mM MgCl_2_, 0.2%, pH 7.2 and 0.2% bovine serum albumin and incubated at 25°C. Oxygen consumption was measured in the homogenates using the Oroboros O2k (Oroboros Instruments Corp) with specific substrates and inhibitors for complex I (pyruvate/proline and rotenone), complex III (glycerol-3-phosphate and antimycin A) and complex IV [ascorbate/TMPD (N,N,N′,N′-tetramethyl-p-phenylenediamine) and cyanide] (Scialo *et al*, 2016a). An unpaired *t*-test with Welch’s correction was used to compare control and RNAi-treated flies.

### Cell culture

HeLa cell lines were cultured at 37°C and 5% CO₂ in high glucose Dulbecco’s modified Eagle’s medium (DMEM; Sigma-Aldrich D5796) supplemented with 10% fetal bovine serum (FBS; Gibco, 17974731), 1× glutaMAX^TM^ (Gibco, 35050061) and 50 μg/ml uridine (Sigma-Aldrich U6375). *PTCD1* cDNAs (wild-type and variants) were synthesized by GenScript (Piscataway, NJ, USA) and cloned into the pBABE-puro vector for retroviral transduction. All constructs were validated by Sanger sequencing. The *PTCD1* knockout cell line (Fleck *et al*., 2019) was a kind gift from Baris Bingol (Department of Neuroscience, Genentech, CA, USA).

### Immunoblotting of cell lines

HeLa variant cells were solubilised in phosphate-buffered saline, 1% DDM, 1 mM PMSF, and complete protease inhibitor (Thermo Fisher Scientific). Protein concentrations were measured by the Bradford assay (Bio-Rad). Equal amounts of proteins were separated by Tris-glycine SDS-PAGE and transferred to nitrocellulose by semi-dry transfer. Membranes were blocked in Tris-buffered saline, 0.1% Tween 20 (TBS-T) with 5% BSA at room temperature for 1 h and primary antibodies were incubated overnight at 4°C in 5% BSA/TBS-T. Membranes were stripped, when required, for 10 min using Pierce™ Restore™ PLUS Western Blot Stripping Buffer (Thermo Fisher Scientific, 46430).

Detection was done the following day using HRP-conjugated secondary antibodies (Jackson ImmunoResearch) and enhanced chemiluminescence (ECL) substrate 20X LumiGLO^®^ Reagent and 20X Peroxide (Cell Signaling Technology 7003). Images are taken on ChemiDoc MP System (Bio-Rad) using Image Lab software. Primary antibodies from Proteintech were used against MRPL11 (15543-1-AP, 1:5000) and MRPS27 (17280-1-AP, 1:5000). The following primary antibodies were obtained from Abcam: PTCD1 (ab121620, 1:2000); OXPHOS Cocktail (ab110411; 1:2000; detecting ATP5A and UQCRC2) and SDHA (ab14715, 1:5000). OPA1 antibody (612606) was obtained from BD Biosciences. Representative immunoblots from independent experiments were processed and assembled using Adobe Photoshop and Illustrator. Linear corrections to brightness were applied uniformly across the images.

### *PTCD1* variant cell line proteomics

Cell pellets were resuspended for 30 min in DDM solubilization buffer containing 1% DDM (Sigma-Aldrich D4641), cOmplete™ EDTA-free protease inhibitor (Roche 64755100), 1 mM PMSF (Sigma-Aldrich P7626), 1 mM dithiothreitol (DTT; Sigma-Aldrich D9779). The suspension was centrifuged at 17 000 g for 20 min, at 4°C to separate soluble and insoluble fractions. Total cellular soluble and insoluble protein fractions were prepared from four biological replicates per variant. The lysis buffer used to solubilize the insoluble pellet fraction was replaced with 50 mM triethylammonium bicarbonate (TEAB; Sigma-Aldrich T7408) at pH 7.55, supplemented with 5% SDS, cOmplete™ Protease Inhibitor Cocktail (Roche), and Pierce™ Universal Nuclease (Thermo Fisher Scientific). Lysates were centrifuged at 16 000 g for 20 min at 4°C to remove debris. Protein concentrations were quantified using the Pierce™ Bicinchoninic Acid (BCA) Protein Assay Kit (Thermo Fisher Scientific). Proteins were reduced with 10 mM tris(2-carboxyethyl)phosphine (TCEP) for 20 min at 37°C and alkylated with 10 mM iodoacetamide for 20 min in the dark. Samples were subsequently digested using S-Trap™ microcolumns (ProtiFi) according to the manufacturer’s instructions. Resulting peptides were dried in a SpeedVac™ vacuum concentrator and stored at –80°C prior to analysis. Dried peptide samples were resuspended in HPLC-grade water containing 0.1% formic acid at 500 ng/μL. Liquid chromatography (LC) was performed using an Evosep One system with a 15 cm Aurora Elite C18 column with an integrated CaptiveSpray emitter (IonOpticks) maintained at 50°C. Buffer A consisted of 0.1 % formic acid in HPLC-grade water and buffer B consisted of 0.1 % formic acid in acetonitrile. Immediately prior to LC–MS analysis, peptides were resuspended in 30 µl of buffer A and a volume corresponding to 500 ng of peptide was loaded onto LC system-specific C18 EvoTips, according to the manufacturer instructions. Samples were analysed using the predefined Whisper-Zoom 20 SPD protocol (0–35 % buffer B gradient at 100 or 200 nl/min for 58 minutes). The Evosep One system was used in line with a timsToF-HT mass spectrometer (Bruker) operated in DIA-PASEF mode. Mass and ion mobility ranges were 300–1200 m/z and 0.6–1.45 1/K0 respectively. diaPASEF aquisition was performed as described previously (Hermosilla-Trespaderne *et al*, 2024) using variable-width ion mobility-m/z windows without overlap. TIMS accumulation and ramp times were each set to 66 ms, resulting in a total cycle time of approximately 1.8 seconds. Collision energy was applied linearly across the ion mobility range (0.6–1.6 1/K0), corresponding to and collision energies of 20–59 eV.

### Retroviral transduction of HeLa cells with *PTCD1* variants

HeLa *PTCD1* knockout cells were transduced with *PTCD1* variants using a pBABE-based retroviral expression system. Retroviral particles were produced in Phoenix-AMPHO packaging cells (CRL-3213™, ATCC). Phoenix-AMPHO cells were cultured in DMEM supplemented with FBS, and uridine under standard conditions (37°C, 5% CO₂) and seeded in six-well plates for transfection. At approximately 50% confluency, Phoenix-AMPHO cells were transfected with pBABE plasmids encoding PTCD1 constructs (WT, R113W, G184R, or R113W/G184R) using jetPRIME transfection reagent (Polyplus) according to the manufacturer’s instructions. Briefly, 2 µg plasmid DNA was diluted in 200 µl jetPRIME buffer, mixed with 4 µl jetPRIME reagent, incubated for 10 min at room temperature, and then added dropwise to the cells. Transfected cells were incubated for 24 h to allow retroviral particle production. Viral supernatants were collected 24 h and 48 h post-transfection, clarified by filtration through 0.45 µm syringe filters, and supplemented with polybrene (final concentration 5 µg/ml) to enhance infection efficiency. HeLa target cells were seeded in six-well plates and infected at 50–70% confluency with viral supernatants containing polybrene. Cells were incubated with viral particles for 24 h, after which the medium was replaced with fresh growth medium. Stable integration of the pBABE constructs was achieved by puromycin selection. Twenty-four hours after infection, puromycin (1.0 µg/ml) was added to the culture medium, and selection was continued for 48–72 h with medium replacement every 24 h to remove dead cells. A non-transduced control well was included to confirm puromycin sensitivity. Once control cells had been eliminated, the surviving transduced cells were expanded for downstream experiments.

### Immunofluorescence staining and microscopy

*PTCD1* and control cells were cultured in DMEM supplemented with 50 µg/ml uridine (Sigma-Aldrich, U3750) and 10% fetal bovine serum (FBS; Gibco, 17974731) at 37°C in a humidified atmosphere containing 5% CO₂. Cells were plated on glass coverslips (ZEISS, thickness 0.17 mm, 22 × 22 mm; 4030-9020-000). After 48 h, cells were rinsed with PBS and fixed in 4% paraformaldehyde (Thermo Scientific, J61899) and 0.02% glutaraldehyde (EM grade; Bangs Laboratories, AA012) in PBS for 15 min at 37°C. Cells were washed three times for 5 min in PBS with gentle rocking and permeabilized with 0.1% Triton X-100 in PBS for 10 min. Samples were then blocked for 60 min at room temperature in PBS containing 5% bovine serum albumin (BSA; IgG-free, protease-free; Jackson ImmunoResearch). Coverslips were incubated overnight at 4°C with primary antibodies diluted in PBS containing 0.1% BSA: mouse anti-ATP5A (Abcam, ab14748; 1:200) and rabbit anti-MRPL11 (Proteintech, 15543; 1:200). After three washes in PBS (5 min each), cells were incubated for 1 h at room temperature in the dark with species-appropriate secondary antibodies: Alexa Fluor 594 goat anti-mouse IgG (Invitrogen, A-11005; 1:200) and Atto 647 goat anti-rabbit IgG (Sigma-Aldrich, 40839; 1:200). Coverslips were mounted using ProLong Glass Antifade Mountant (Invitrogen, REFP36980; refractive index 1.52) and allowed to cure for 24 h before imaging. Images were acquired using an Olympus FV4000 confocal microscope equipped with a UPLXAPO 60×/1.42 NA oil immersion objective. Fluorophores were excited using 594- and 640-nm laser lines. Z-stacks were collected with a step size of 0.12 µm and a pinhole setting of 0.8 Airy units. Image processing and analysis were performed using Fiji/ImageJ. Only linear adjustments were applied and were identical across all images. Mitochondrial fragmentation was quantified from n > 30 cells per condition across three independent experiments.

### Quantification and statistical analysis

Raw data files were initially searched using DIA-NN 2.0.1 to generate an in silico spectral library from the *Homo sapiens* (downloaded from Uniprot on the 15th of August 2023, containing 20 423 entries) proteome, as well as a common contaminants list. This library was generated in DIA-NN 2.0.1 (Demichev *et al*, 2020) with fixed carbamidomethyl (C) and variable oxidation (M) peptide mass modifications, and mass accuracy and MS1 accuracy set to 15. Peptide length range was 7–30, precursor charge range 1–4, precursor m/z range 300–1800, and fragment ion m/z range 200–1800. The search output was filtered to remove common contaminants, proteins only identified by one unique peptide, and any proteins not found in at least three replicates of one group. The data was then log_2_ transformed, normalised based on median intensities, followed by limma analysis (Ritchie *et al*, 2015) in RStudio to calculate fold changes and *p* values. Volcano plots were then generated based on this data.

Mitochondrial fragment size values were normalized and visualised as violin plots, including all measurements pooled across experiments without separation by replicate. Fragment sizes were analysed over a range of ≥ 0.2 µm², representing the full distribution of detected mitochondrial fragments. Each data point represents individual mitochondrial fragment. Statistical significance between groups was assessed using the two-sample Kolmogorov–Smirnov (KS) test to compare the distributions of mitochondrial fragment sizes between conditions.

For the cardiac mitochondrial proteomics experiment, raw files were processed using MaxQuant (v.1.6.0.16) (Cox & Mann, 2008) and data analysis performed using Perseus (Tyanova *et al*., 2016). For detailed instrument parameters and MaxQuant search settings, see (Hock *et al*., 2020). ProteinGroups file was loaded onto Perseus (v.1.6.14.0) and proteins annotated for contaminants, only identified by site reverse, and identified by single peptide were removed from the dataset. Relative Complex Abundance (RCA) plot was generated using the RCA tool available on www.rdms.app with significance calculated from a two-sided t-test between the individual protein means (Hock *et al*., 2025). MitoCarta3.0 annotation (Rath *et al*., 2021) was used to subset mitochondrial proteins and two valid values in each group (patient or controls) prior to mean normalisation and two-sample, two-tailed Student’s *t*-test to generate the volcano plots. Topographical heatmap was generated using the log_2_ fold changes from *t*-test as previously (Stroud *et al*, 2016) using the structure of the human mitochondrial ribosome PDB ID: 3J9M (Amunts *et al*., 2015).

## Data availability

The heart mitochondria and cell lines mass spectrometry proteomics data have been deposited to the ProteomeXchange Consortium via the PRIDE partner repository (Perez-Riverol *et al*, 2025) with the dataset identifier PXD078959 and PXD079210, respectively. The source data of this paper are collected in the following database record.xxx

## Acknowledgements

M.O is supported by Fight for Sight and The Academy of Medical Sciences. This research was funded by the Jane and Aatos Erkko Foundation, the Sigrid Jusélius Foundation, the Research Council of Finland, and the Newcastle University Academic Track (NUAcT) Fellowship awarded to U.R. R.W.T is funded by the Wellcome Centre for Mitochondrial Research (203105/Z/16/Z), the Medical Research Council (MR/W019027/1), the UK NIHR Biomedical Research Centre for Ageing and Age-related disease award to the Newcastle upon Tyne Foundation Hospitals NHS Trust, the Lily Foundation, the LifeArc Centre for Rare Mitochondrial Diseases and the UK NHS Highly Specialised Service for Rare Mitochondrial Disorders of Adults and Children. A.A acknowledges support from the Finnish Cultural Foundation and the Jenny and Antti Wihuri Foundation through the Postdoc Pooli Grant Scheme, Finland. A.A, N.W and J.M are funded by the Jane and Aatos Erkko Foundation (240022). A.S was supported by two BBSRC grants (BB/R008167/2 & BB/W006774/1) and a Wellcome Senior Research Fellowship (212241/A/18/Z). This project was also supported by grants from the Australian National Health and Medical Research Council (NHMRC) to D.A.S (2009732), the Medical Research Futures Fund (MRFF) to D.A.S and D.H.H (MRF2016030), MRFF National Critical Research Infrastructure program to D.A.S and D.H.H (NCRI000043), and the Mito Foundation research equipment grants to D.A.S and D.H.H (G189). We thank Dr Ashwin Sriram for his help with aspects of the project involving the use of *Drosophila melanogaster*. We thank Dr Charlotte Alston for her assistance with variant interpretation and allele frequency analysis. We thank Margaux Hedin for her work on mitochondrial stress signalling. Imaging was performed at the Light Microscopy Unit, Institute of Biotechnology, supported by HiLIFE and Biocenter Finland.

## Expanded view figures

**Figure EV1.** Molecular cloning and subsequent Sanger sequencing of individual cDNA clones was used to establish the segregation pattern of *PTCD1* variants. (**A**) Table showing each variant position in seven Sanger sequenced *PTCD1* patient cDNA clones compared to the reference sequence (NM_015545.4). The c.337C>T, p.(Arg113Trp) and c.550G>A, p.(Gly184Arg) missense variants segregate together in *cis*, whilst the c.388C>T, p.(Arg130*) truncating variant is present on the other allele. (**B**) Summary of *PTCD1* allelic phasing indicated the allelic configuration and predicted functional effect.

**Figure EV2. Predicted 3D structures of *PTCD1* variants.** AlphaFold models generated using ColabFold are shown for wild-type PTCD1 and the indicated single and double amino acid substitution variants. Residue numbers denote positions within the PTCD1 polypeptide, with the original residue indicated to the left and the substituted residue to the right of the position number. Variant residues are highlighted and indicated by white arrows, and a key structural element of the protein is enlarged for clarity. The R113W variant is located within an unstructured domain, typical for RNA–protein chaperones, whereas the G184R variant lies within the RNA-interacting PPR domain of PTCD1. Colours are used solely to distinguish the different variants and do not imply any biological or functional differences.

**Figure EV3.** (**A**) PTCD1-IR male flies 1 day post eclosion compared to control flies. The genotypes used were as follows: Ctrl = w1118;daGAL4, PTCD1-IR = w1118;PTCD1-IR;daGAL4. (**B**) Respirometry measurements of control and PTCD1-IR male whole-fly homogenates were performed using an OROBOROS O2k oxygraph. CI-linked respiration (CI+CIII+CIV) was initiated by the addition of 5 mM pyruvate, 5 mM proline and 1 mM ADP and then inhibited by adding 0.5 μM rotenone. CIII-linked respiration (CIII+CIV) was stimulated by the addition of 20 mM glycerol 3-phosphate and then inhibited with the addition of 2.5 μM antimycin A. CIV respiration (CIV) was initiated by the addition of 4 mM ascorbate and 2 mM TMPD and inhibited by adding 0.5 mM KCN. Statistical analysis using a *t*-test with Welch’s correction was performed (n = 8–9 samples per group).

**Figure EV4. Immunoblot analysis of mitochondria and post-nuclear supernatant isolated from control and PTCD1 patient cardiac tissues**. Samples were lysed in the presence of 1% SDC and heated at 99°C, for 15 min to solubilise both soluble and insoluble protein fractions, that were subsequently used for proteomics analysis. Antibodies against PTCD1, mtLSU MRPL28, mtSSU MRPS17, OXPHOS complex I NDUFB8, complex II SDHB subunits, FIS1 and mtHSP70 were used. Due to limited availability of patient cardiac tissue, n = 1. Mito, mitochondrial fraction, PNS, post-nuclear supernatant.

**Figure EV5. Biological replicates for mitochondrial fragment size. (A)** Violin plot of *PTCD*1 control, variants, and empty vector transduced *PTCD1* knockout HeLa cells illustrating fragment size. No significant differences were observed between the two single variants and the double variant, whereas both the control and the PTCD1 knockout differed significantly from all other conditions. The KS test was performed on average mitochondrial fragment size per cell to visualise significance with corresponding p-values (*p = 0.05, **p = 0.01, ***p = 0.001, ****p = 0.0001, NS = p > 0.05, n = 116). Graphs are accompanied by representative confocal images of ATP5A-stained cells for each condition. Acquired on Olympus FV4000, UPLXAPO 60x/1.42 oil WD 0.15 mm. **(B)** Biological replicates for mitochondrial fragment size. Violin plots showing the distribution of normalised mitochondrial fragment area (um²) across three biological replicates within each condition and between conditions. Colours were used for visual distinction only. **(C)** Gaussian mixture model-based classification of fragmentation states. Data from all variant transductions were used to define four thresholds separating distinct degrees of mitochondrial fragmentation, ensuring consistent classification across conditions. Thresholds were derived from the intersections between Gaussian components. These discrete categories were used to group data for bar graph analyses.

**Fig. EV6.** A summary allele phasing indicating that PTCD1 patient is carrying a hypomorphic complex allele along with a loss-of-function allele, with *PTCD1* variants p.(Arg113Trp); p.(Gly184Arg) in *cis* with p.(Arg130*) in *trans*.

