## Supplemental materials (EV) for "Pathogenic *PTCD1* variants cause mitochondrial protein aggregation and cardiomyopathy": FiguresEV.pdf

# EV1

## A

| | Reference variant NM_015545.4 | Clone 1 | Clone 2 | Clone 3 | Clone 4 | Clone 5 | Clone 6 | Clone 7 |
|---|---|---|---|---|---|---|---|---|
| c.337C>T, pArg113Trp | G | **T** | **T** | **T** | G | **T** | **T** | **T** |
| c.388C>T, pArg130* | G | G | G | G | **A** | G | N | G |
| c.550G>A, pGly184Arg | C | **A** | **A** | **A** | C | **A** | **A** | **A** |

## EV2

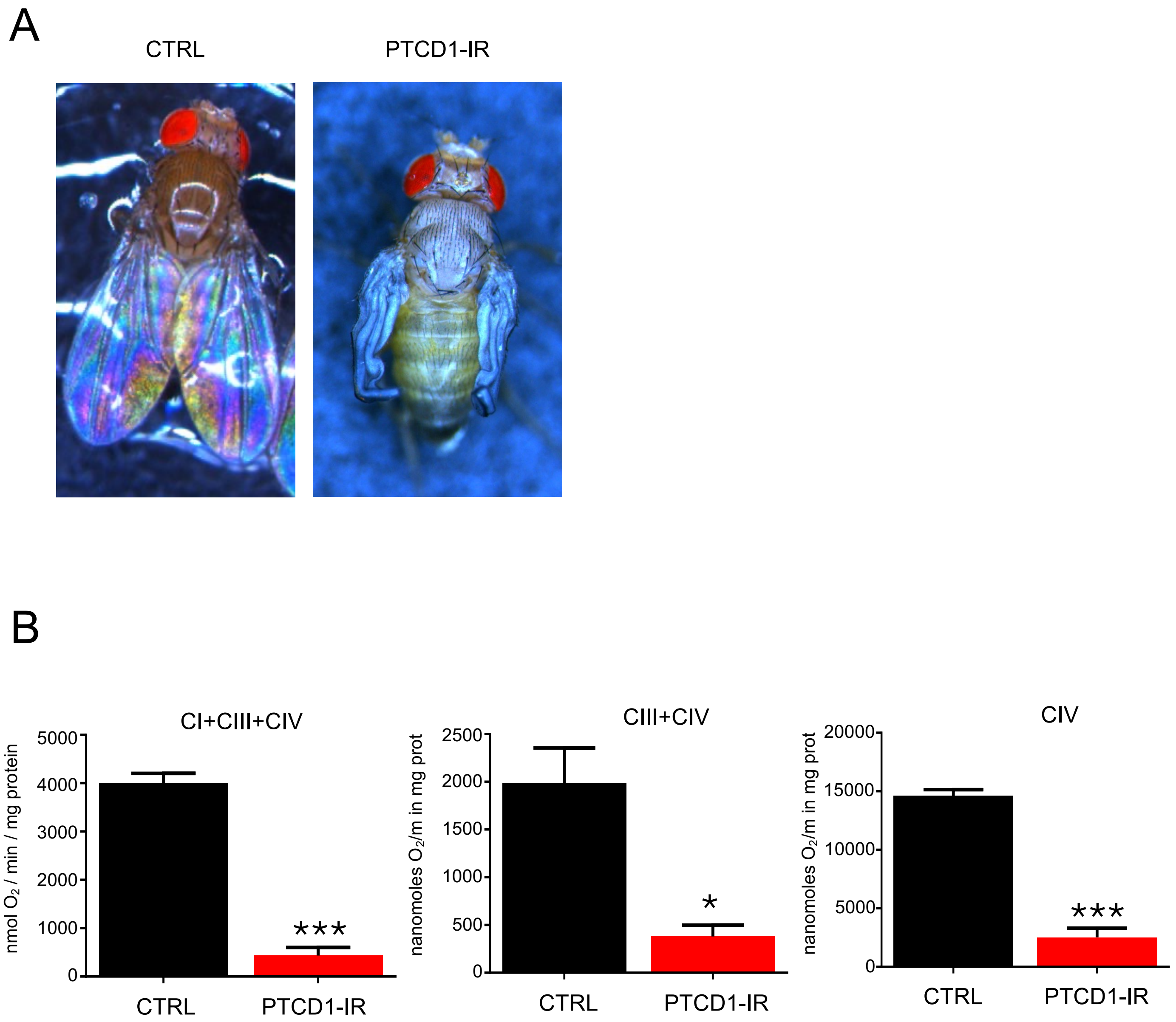
A
CTRL
PTCD1-IR
B
CI+CIII+CIV
nmol O2 / min / mg protein
5000
4000
3000
2000
1000
0
***
CTRL
PTCD1-IR
CIII+CIV
nanomoles O2/m in mg prot
2500
2000
1500
1000
500
0
*
CTRL
PTCD1-IR
CIV
nanomoles O2/m in mg prot
20000
15000
10000
5000
0
***
CTRL
PTCD1-IR

# EV 3

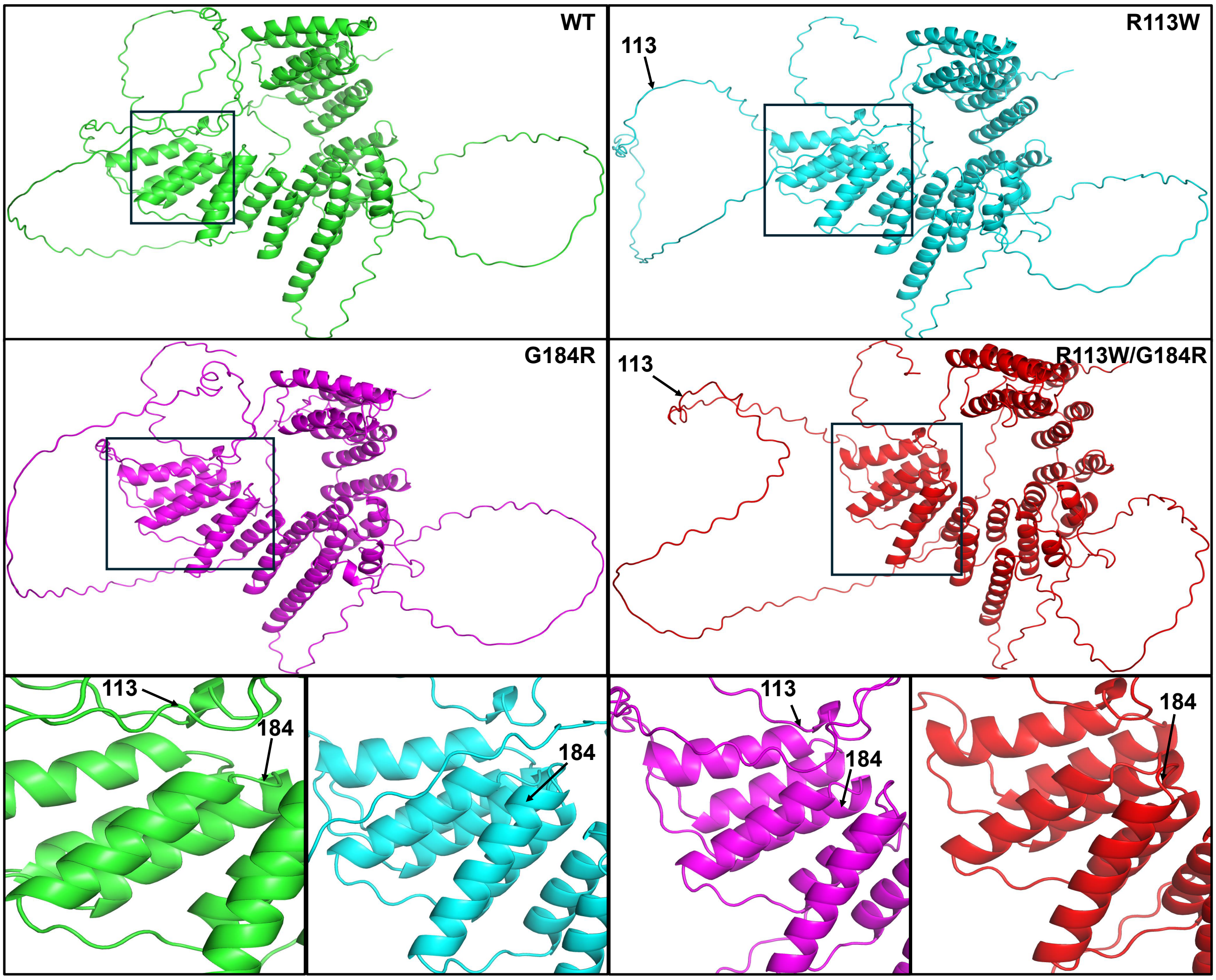

# EV 4

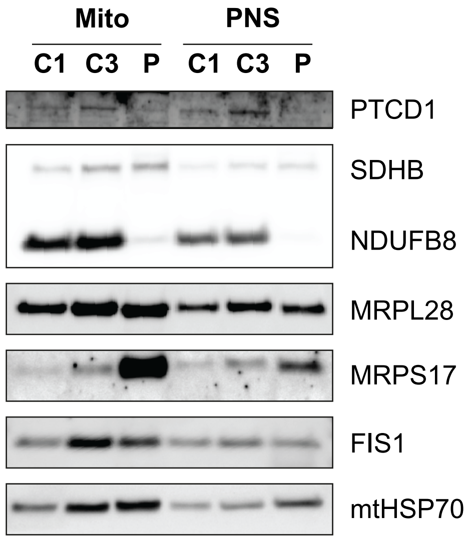
Mito
PNS
C1
C3
P
C1
C3
P
PTCD1
SDHB
NDUFB8
MRPL28
MRPS17
FIS1
mtHSP70

## EV 5

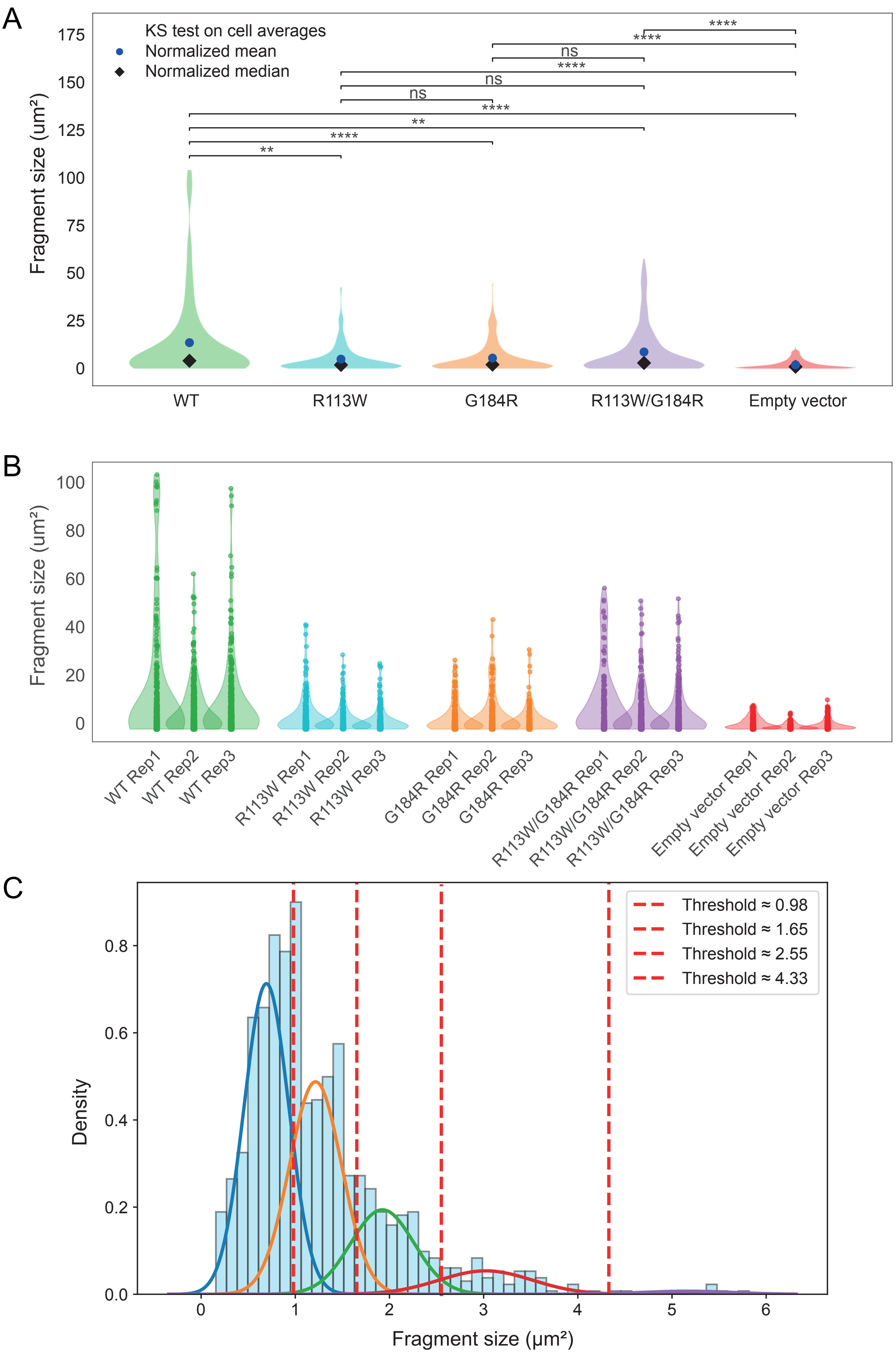
A
KS test on cell averages
Normalized mean
Normalized median
Fragment size (um²)
175
150
125
100
75
50
25
0
ns
ns
ns
****
**
**
WT
R113W
G184R
R113W/G184R
Empty vector
B
Fragment size (um²)
100
80
60
40
20
0
WT Rep1
WT Rep2
WT Rep3
R113W Rep1
R113W Rep2
R113W Rep3
G184R Rep1
G184R Rep2
G184R Rep3
R113W/G184R Rep1
R113W/G184R Rep2
R113W/G184R Rep3
Empty vector Rep1
Empty vector Rep2
Empty vector Rep3
C
Threshold ≈ 0.98
Threshold ≈ 1.65
Threshold ≈ 2.55
Threshold ≈ 4.33
Density
0.8
0.6
0.4
0.2
0.0
0
1
2
3
4
5
6
Fragment size (µm²)

# EV 6

| Variant ID | cDNA change | Protein change | Shorthand | Variant type | Allelic configuration | Predicted functional effect |
|---|---|---|---|---|---|---|
| Variant A | c.337C>T | p.(Arg113Trp) | R113W | Missense | In *cis* with p.(Gly184Arg) | Likely hypomorphic |
| Variant B | c.550G>A | p.(Gly184Arg) | G184R | Missense | In *cis* with p.(Arg113Trp) | Likely hypomorphic |
| Variant C | c.388C>T | p.(Arg130*) | R130* | Nonsense / truncating | In *trans* | Predicted loss-of-function |
| Complex allele | c.337C>T + c.550G>A | p.(Arg113Trp);p.(Gly184Arg) | R113W/G184R (dual) | Complex missense allele | In *cis* | Likely hypomorphic complex allele |
